# Comprehensive cell atlas of the first-trimester developing human brain

**DOI:** 10.1101/2022.10.24.513487

**Authors:** Emelie Braun, Miri Danan-Gotthold, Lars E. Borm, Elin Vinsland, Ka Wai Lee, Peter Lönnerberg, Lijuan Hu, Xiaofei Li, Xiaoling He, Žaneta Andrusivová, Joakim Lundeberg, Ernest Arenas, Roger A. Barker, Erik Sundström, Sten Linnarsson

## Abstract

The adult human brain likely comprises more than a thousand kinds of neurons, and an unknown number of glial cell types, but how cellular diversity arises during early brain development is not known. Here, in order to reveal the precise sequence of events during early brain development, we used single-cell RNA sequencing and spatial transcriptomics to uncover cell states and trajectories in human brains at 5 – 14 post-conceptional weeks (p.c.w.). We identified twelve major classes and over 600 distinct cell states, which mapped to precise spatial anatomical domains at 5 p.c.w. We uncovered detailed differentiation trajectories of the human forebrain, and a surprisingly large number of region-specific glioblasts maturing into distinct pre-astrocytes and pre-oligodendrocyte precursor cells (pre-OPCs). Our findings reveal the emergence of cell types during the critical first trimester of human brain development.

---

The cellular architecture and wiring of the human brain have evolved to support the most complex behaviors known in all biology, such as integrated sensorimotor skills, language, and cognitive abilities that surpass those of any other animals. Like all mammalian brains, the human brain develops by initial patterning of the neural tube, through a series of specification, differentiation and maturation events that yield likely more than a thousand distinct types of neurons, as well as glia and non-neural cell types. The fundamental architecture of the human brain follows from these patterning events, and can be described using the prosomeric model, which divides the brain into antero-posterior neuromeres and dorso-ventral plates (floor, basal, alar and roof plates) (1). While less is known about regional patterning of macroglia, emerging evidence from the mouse suggests that astrocytes can be divided into major telencephalic and non-telencephalic types, whereas oligodendrocytes appear to be transcriptionally less heterogeneous throughout the brain. Single-cell transcriptomics has emerged as a powerful approach to reveal cell types and differentiation trajectories during development. A number of studies have explored specific regions during human brain development, particularly the neocortex (2–6). However, no comprehensive study exists of the whole human brain during the crucial first trimester when the brain is patterned.

## A census of developmental cell states

Here we describe the developing human brain using RNA sequencing of single cells from 26 dissected embryonic and fetal brains spanning 5 to 14 weeks post-conception (p.c.w.; table S1 and fig. S1). 15 donors were female based on *XIST* expression (fig. S1A). We collected 111 unique biological samples (340 including technical replicates; fig. S1F and table S1) and sequenced more than two million cells using droplet-based single-cell RNA sequencing. Two versions of the 10X Chromium single-cell RNA-seq chemistry were used (v2 and v3), resulting in an average of 4,262 and 10,940 UMIs per cell, respectively (fig. S1A-E). We corrected for batch effects caused by chemistry differences using Harmony (7), resulting in a well-integrated dataset (fig. S1C-D). We used stringent automated quality control (fig. S2A-F) to reject empty droplets, cellular debris, bare nuclei, low-quality cells and doublets, followed by manual curation resulting in 1,665,937 high-quality cells. We used a conventional analysis pipeline (Methods) to define 616 robust clusters, and annotated those clusters with metadata including class and subclass, spatial location, embryonic age distribution, and specific markers (table S2). Since regional dissections of human embryos is challenging, we validated the spatial location of each cell by training and applying a classifier, thus inferring the provenance of cells in all samples from their expression profiles. Reassuringly, samples dissected as whole brain showed proportions of cells from forebrain, midbrain and hindbrain, whereas samples dissected from single regions showed cells predominantly from that region (fig. S2H).

The dataset was dominated by cells of the neuronal lineage (Fig. 1A-C): radial glia (defined by expression of *HES1*), neuroblasts (*NHLH1*), neuronal intermediate progenitor cells (IPCs; defined as neuroblasts and immature neurons expressing cell cycle genes), and immature neurons (expressing the neurofilament gene *INA*). At later timepoints, and in the direction away from neurons, radial glia matured into putative glioblasts (defined here by expression of *TNC* and *BCAN*) and oligodendrocyte precursor cells (OPCs; expressing *PDGFRA* and *OLIG1*). Smaller populations of cells, originating from outside the neural tube proper, included vascular and blood cells (endothelial cells, pericytes, erythrocytes), immune cells (microglia), mesenchymal fibroblasts, and cells derived from the placodes and the neural crest. Quantifying the abundance of major cell classes by developmental timepoint (Fig. 1B, adjusted for sampling depth and brain volume; see Methods), neurons and neuroblasts showed relatively constant abundances. Neuronal IPCs increased by age, as did cells of the immune, oligodendrocyte and vascular lineages. Instead, the major change over the timespan studied was the radial glia to glioblast switch, which occurred around 10.5, 9.5 and 7.5 p.c.w. in forebrain, midbrain and hindbrain, respectively.

**Figure 1.**
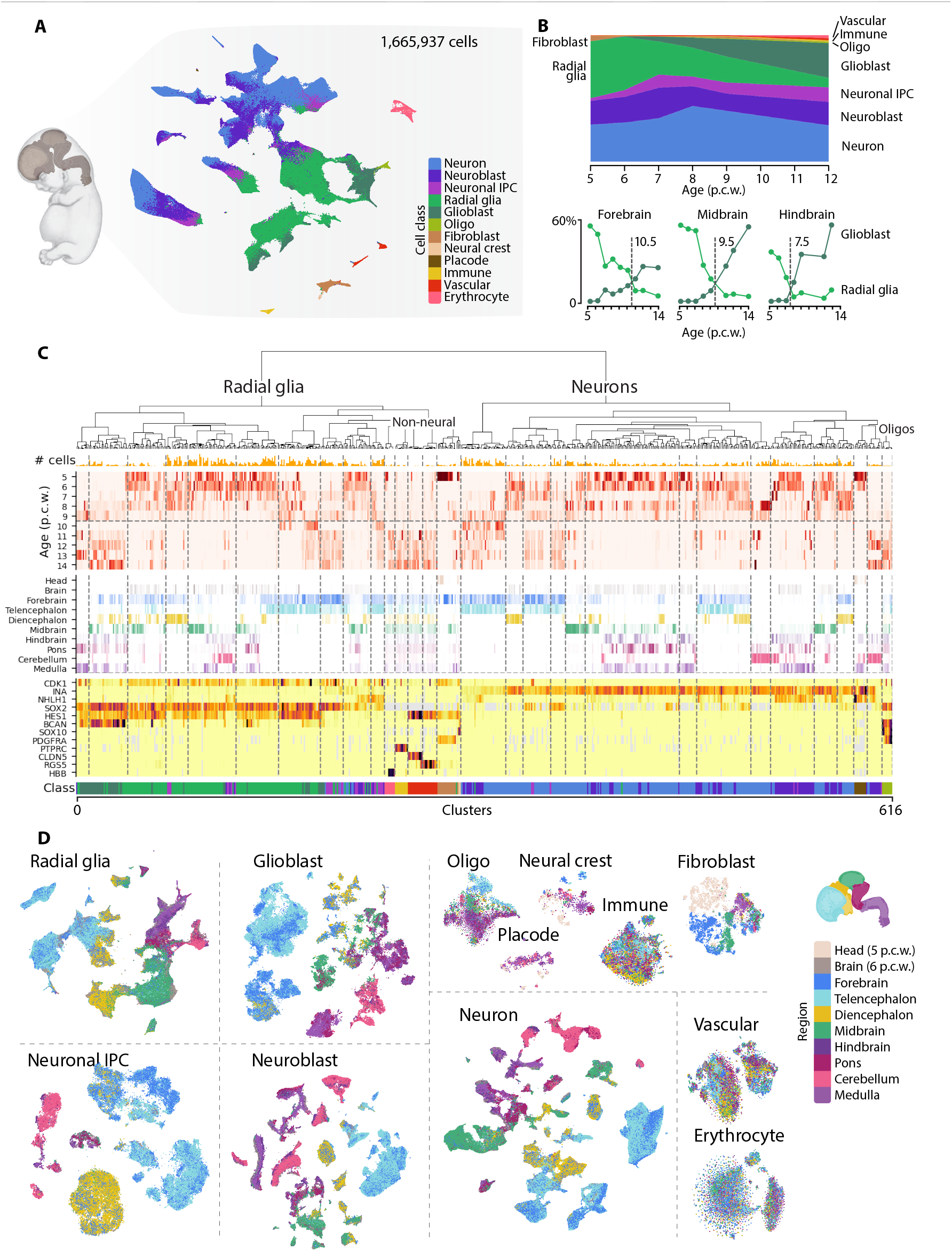
Overview. **(A)** tSNE-plot showing major cell classes. **(B)** Top, proportion of cell classes by age, omitting 13 and 14 p.c.w. because of incomplete sampling at those ages. Bottom, proportion of glioblasts and radial glia by developmental age and brain region. **(C)** Metadata for all 617 clusters, including age, regional distribution, common cell type markers and class (colored as in A). **(D)** tSNE plots showing the regional identity of cells belonging to each major class. Embryo image in (A) used with permission from HDBR Atlas (https://hdbratlas.org).

The fundamental architecture of the human brain is established by patterning of the neural tube along antero-posterior and dorso-ventral axes, onto which is superimposed the maturation of radial glia into neurons along the radial direction. As expected, we found that radial glia, neuronal IPCs, neuroblasts and neurons all showed strong regional patterning (Fig1D). We further found a rich repertoire of region-specific glioblasts, with distinct subtypes corresponding to each developmental compartment (Fig 1D). OPCs were also patterned along the antero-posterior axis (see further below). In contrast, immune, vascular and blood cells lacked strong region-specificity. In order to learn the detailed spatial location of cell types during the early patterning phase, we performed multiplexed RNA fluorescent *in situ* hybridization (EEL FISH (*8*)) targeting 440 selected genes in three relatively medial sagittal sections of a single embryo at postconception week 5 (Fig. 2, fig. S4, and table S3), complemented by transcriptome-wide spatial transcriptomics (10X Visium) on eight sections sampled more widely (fig. S4E). We used the observed spatial expression of transcription factors and morphogens with known anatomical expression patterns (Fig. 2A and fig. S4B) to manually curate an anatomical atlas of the specimen following the prosomeric model (Fig. 2B and fig. S4A). We further used the expression of maturation markers (*SOX2*, radial glia; *NHLH1*, early neuroblasts; *NRXN3*, late GABAergic neuroblasts; *STMN2*, neurons) to designate the ventricular, subventricular and mantle zones along the extent of the neural tube (fig. 2B, S4C and S4E). Next, we aligned scRNA-seq clusters with spatial regions using BoneFight (*9, 10*), considering only those clusters with sufficient abundance at 5 p.c.w. (Methods). This resulted in the alignment of 64 clusters, and revealed highly detailed spatially restricted domains consisting of transcriptionally defined cell types (Fig. 2C and fig. S4D).

**Figure 2.**
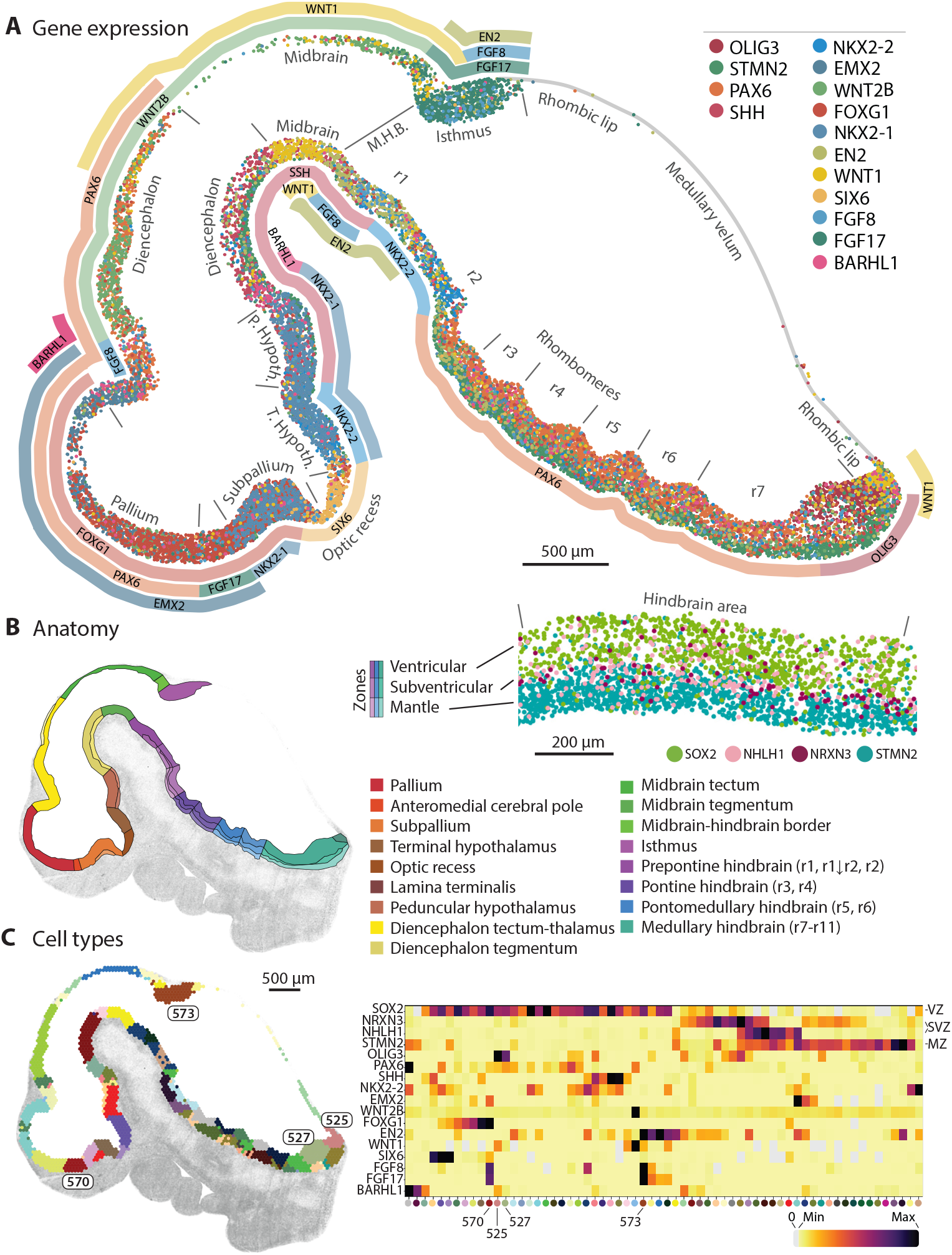
Spatial gene expression at 5 p.c.w. **(A)** Scatter plot showing mRNA molecules detected in the neural tube of a single sagittal section (molecules outside the neural tube are not shown), colored by gene identity. Prosomeres are indicated and approximate gene expression domains are shown as ribbons. **(B)** Manually curated anatomical domains following the prosomeric model. Inset, expression of markers of the ventricular, subventricular and mantle zones shown for a segment of the ventral hindbrain. **(C)** Left, single-cell clusters assigned to spatial domains using BoneFight. Right, heatmap showing the expression of the same genes as in (A) across aligned clusters.

## Excitatory lineage of the developing neocortex

The cerebral cortex contains two main classes of neurons, glutamatergic excitatory neurons and GABA (γ-aminobutyric acid)-ergic inhibitory interneurons. Excitatory neurons are generated by progenitors located in the developing cortex (i.e., the pallium) and migrate radially. In contrast, cortical inhibitory neurons are generated by progenitors in the ganglionic eminences of the ventral forebrain and migrate tangentially into the developing neocortex (*11–13*).

To examine the developmental lineage of the excitatory neurons, we computationally isolated those cortical clusters expressing *EMX1*, a transcription factor that marks the cortical excitatory lineage (*12*). Preliminary analysis revealed clusters that were driven by the precise age, subregion, or donor, and we observed very clear batch differences even beyond technical batch effects due to single-cell chemistry differences. Such strong sample-to-sample differences were not observed in the interneuron lineage, and may be a result of a greater sensitivity of excitatory neurons to genetic differences, or to greater differences in maturation phase or spatial location in the excitatory neuronal lineage.

In order to overcome these batch related differences, we used Scanorama (*14*) to integrate cells from the same post-conceptional week and scVI (*15*) to integrate cells from all time points. The joint embedding revealed a single main lineage with both cell class and cell cycle phase clearly identifiable, and separating a main excitatory neuron lineage from a minor lineage of the *RELN*-expressing Cajal Retzius cells (Fig 3A,D, fig. S5A-D). Radial glia and IPCs each formed loops on the two-dimensional embedding, corresponding to the phases of the cell cycle. Superimposed on each cell cycle loop was an inside-out maturation gradient with older cells located at the outside of each cell cycle loop (Fig. 3H). Thus, the integrated embedding revealed all the key stages of excitatory neuron development: proliferation of radial glia progenitors, differentiation into IPCs, proliferation of IPCs, differentiation of IPCs into neuroblasts, and gradual maturation of neuroblasts into more mature neurons.

**Figure 3.**
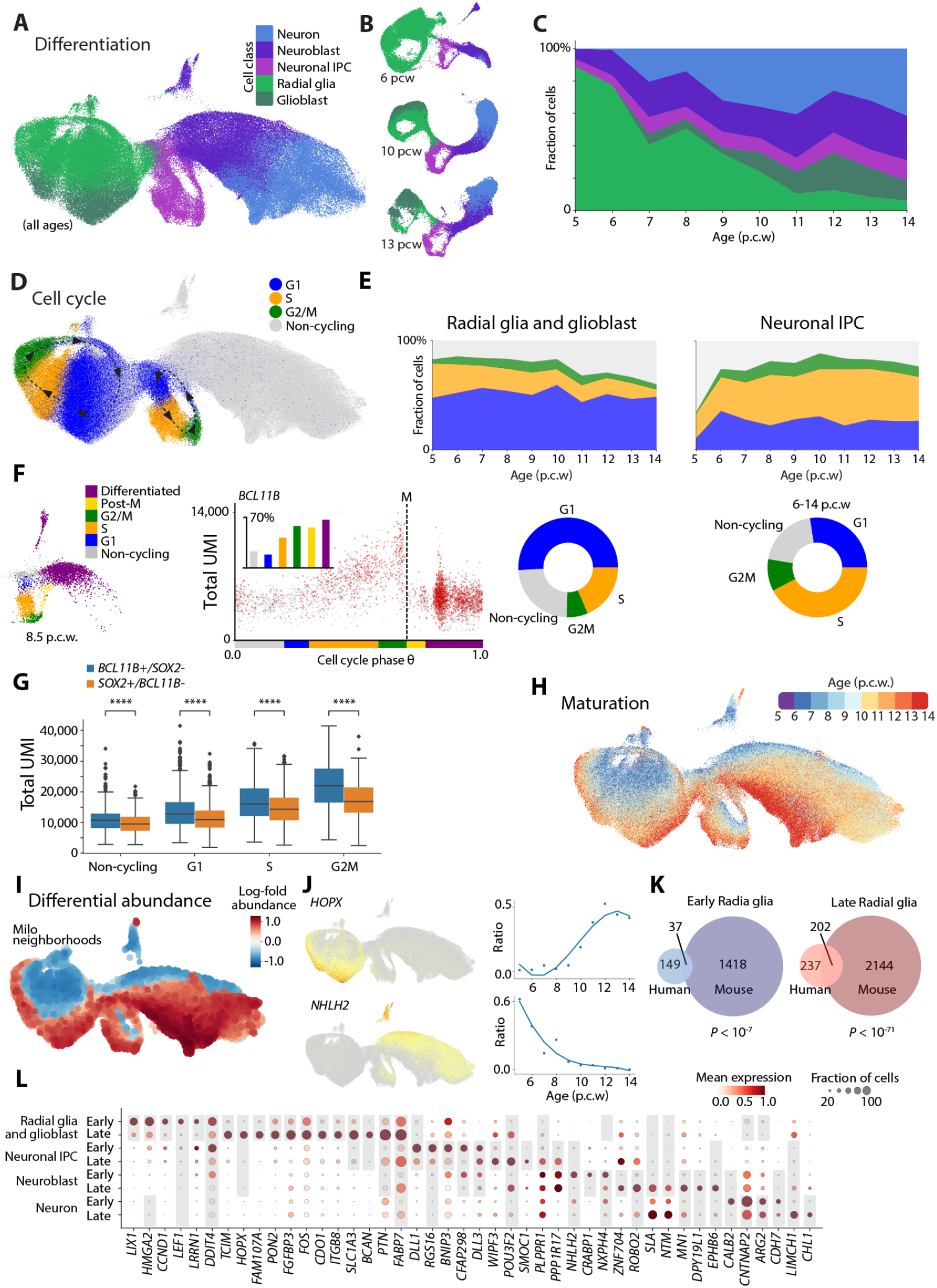
The developing neocortex – excitatory neurons lineage. **(A)** UMAP projections from all collected pallial excitatory neuron lineage (*EMX1*+ cells) colored by major cell classes **(B)** UMAP projections from selected post-conceptional ages (6,9,13 p.c.w) pallium cells (*EMX1*+ cells) colored by major cell classes. **(C)** Distribution of major classes of cells by post-conceptional age. **(D)** UMAP projections from all collected pallial excitatory neuron lineage (*EMX1*+ cells) colored by progenitor state. **(E)** Distribution of progenitor states by post-conceptional age in radial glia cells (left) and neuronal IPC (right). Donut plots show mean values for radial glia in 5-14 p.c.w and neuronal IPCs. **(F)** Box plots showing UMI counts in *BCL11B+/SOX2-vs. SOX2+/BCL11B-*cells by progenitor states in neuronal IPCs in Chromium version 3 samples. **(G)** Progenitor states for neuronal IPCs in 8.5 p.c.w inferred by DeepCycle transcriptional phase (left). UMI counts per cell as a function of the transcriptional phase show growth trend followed by a drop in RNA counts (UMI) that identifies mitosis (marked by M). Cells expressing *BCL11B* are colored red. Small bar plot shows the ratio of cells in each progenitor state expressing *BCL11B* (right). **(H)** UMAP projections from all collected pallial excitatory neuron lineage (*EMX1*+ cells) colored by post-conceptional age. **(I)** Differential abundance across post-conceptional age in pallial excitatory neuron lineage. Milo neighborhood embedding of excitatory neuron lineage. Each point represents a neighborhood, the size of points is proportional to the number of cells in the neighborhood. Neighborhoods are colored by their log-fold change in abundance between early and late post-conceptional age. Neighborhoods showing significant enrichment (SpatialFDR<10%) are colored. Neighborhoods with higher abundance of cells from early ages are colored in blue and neighborhoods with higher abundance of cells from later ages are colored in red. **(J)** Examples of early and late related genes; *HOPX* in late RG and IPCs and *NHLH2* in early neuroblasts and neurons. Cells colored by gene expression (left) Ratio of gene expressing cells in every post-conceptional week (right). **(K)** Overlap of human and mouse early RG associated genes (left) and late associated genes (right). **(L)** Differential expressed genes between early and late states. Selected genes for every class are shown. Dotplot showing mean expression (color) and ratio of cells expressing the gene (circle size) in each class early and late cells.

To confirm that the integrated embedding accurately depicted the developing tissue at all ages, we examined samples from each postconceptional week separately (Fig. 3B, fig. S5B). Radial glia, IPCs and neuroblasts were detected already at 5 p.c.w, while neurons and the first glioblasts were detected only from 6 p.c.w. As expected, the fraction of neurons and glioblasts out of overall cells increased at later time points whereas the fraction of radial glia progenitors decreased (Fig 3B,C, fig. S5B).

Closer examination of gene expression differences between neurons and neuroblasts showed that genes critical for neuronal migration in the mouse were induced in the neuron class cells, including *DAB1, VLDLR* and *DSCAM* (*16, 17*). In contrast, genes that are uniquely expressed in the germinal zone, such as *PPP17R1* and *NEUROD4* were observed in neuroblasts (*18, 19*) (table S4). Thus, this indicates that the transcriptionally defined neuronal class in the excitatory neuronal lineage corresponds to the migrating or post-migratory neurons while the neuroblasts correspond to newborn neurons that reside in the germinal zone. As noted above, the neuroblast marker *NHLH1* was spatially located to the subventricular zone, sandwiched between the ventricular and mantle zones.

As noted, radial glia/glioblasts and IPCs formed loops ordered by the progression through the cell cycle phases (G1, S, G2, M) (Fig 3D). The proportion of cells in each phase was markedly different between radial glia and IPC progenitors, with most cycling radial glia cells found in the G1 phase while most cycling IPCs were found in S phase (Fig 3E). This reveals two distinct cell cycle dynamics between radial glia and IPC progenitors. Assuming equal cell cycle duration, the findings would indicate differences in the rate of progression through the individual cell cycle stages. Using the magnitude of RNA velocity (i.e. the rate of change of gene expression) we compared the cell cycle length of radial glia and IPCs and found it to be comparable in early time points. However, later radial glia/glioblasts had higher magnitude of RNA velocity as compared to IPCs indicating their cell cycle is shorter (fig. S5E). This is consistent with higher proliferative capacity of late radial glia, mainly in the outer SVZ, observed in species with an expanded neocortex including human (*20–22*). Thus, our results suggest that IPCs have a longer S phase compared to radial glia and glioblasts and in later time points their cell cycle is also longer compared to radial glia/glioblasts.

Pebworth et al. identified two transcriptional subtypes of IPCs; radial glia-like cells that express *SOX2* and neuron-like cells that express neuronal genes like *NEUROD6* (*18*). To explore these subtypes in cortical IPCs, we resolved the cell cycle trajectory for IPCs and early differentiating IPCs in samples from two donors (8.5 and 14 p.c.w) using DeepCycle (*23*), yielding a projection of each cell to an offset along the cell cycle. As predicted, RNA molecule counts (UMIs) increased gradually along the cell cycle, with a sudden drop at mitosis, corresponding to cell growth followed by cell division (Fig. 3F, fig S5F-I). The ratio of cells expressing the neuronal genes *BCL11B, NEUROD6* and *INA* consistently increased as the cell cycle progressed, with a very low expression ratio detected at early G1 phase and the highest ratio in post-mitotic differentiating cells. The radial glia-marker *SOX2* showed the opposite trend with higher ratio of cells expressing the gene during G1 and S phases, and lower ratio at G2/M phases and after differentiation (fig. S5G,I). These patterns of expression were consistent at all examined time points (fig. S5J), and corroborate previous study that showed that the IPC cell-fate decision-point occurs at late G1 phase (*24*).

To shed light on the choice between differentiation and self-renewal of IPCs, we contrasted IPCs expressing neuronal genes (*BCL11B, NEUROD6* and *INA*) but not *SOX2* versus those showing the opposite pattern of expression. Interestingly, we found that the neuronal-like cells had significantly higher RNA molecule counts throughout the cell cycle in most of the examined conditions (Fig. 3G, fig. S5K). This indicates that neuronal-like IPCs express more total RNA and potentially a larger number of genes than radial glia-like IPCs, providing a plausible mechanism linking cell growth to differentiation.

To discover further differences between neuronal-like and radial glia-like IPCs, we compared gene expression to find a subset of radial glia and differentiation-associated genes, and then used those genes to further detect differentially expressed genes between IPCs that enter the cell cycle versus differentiating IPCs. We found a total of 72 genes, with most showing high expression in neuroblast and neurons indicating that they have a role in differentiation and neuron formation (table S5, fig. S5L-N). These genes included *INA, STMN2, SNAP25*, and *BCL11B* that are known neuronal markers. The observation that neuronal genes are starting to be expressed after the G1 phase is consistent with previous studies in mouse that showed that the extended length of G1 phase is critical for the cell to differentiate, and with the “cell cycle length model” that posits that this lengthening of G1 is required for accumulation and/ or action of factors that drive differentiation (*25*). Our results further show that for many neuronal genes, the expression increased gradually across the IPC cell cycle and later along the differentiation axis, starting at IPCs late G1 phase (fig S5M-N).

Although the fate-decision point was observed at late G1 phase, in our resolved cell cycle we observed that differentiation point for both radial glia and IPCs occurred shortly after the M phase. At all ages, and in the integrated embedding, we observed short trajectories bridging radial glia to IPCs and IPCs to differentiating neurons, in each case linking late or post-M phase of a previous stage to the beginning of the next stage (Fig. 3A,D, fig. S5B). In contrast, few if any cells were found to transition from radial glia to IPCs during G1, S or G2/M phases (as indicated by the absence of cells bridging those phases between the two cell types). Furthermore, non-cycling IPCs may plausibly correspond to those radial glia that differentiate directly into neurons. Thus, progenitors decide just after dividing whether to commit to another division, or progress to the next stage of differentiation.

Taken together, these findings support a model in which the vast majority of IPCs start the cell cycle as radial glia-like IPCs. At late G1 phase or early S phase the fate decision is made and the cell proceeds through the cell cycle as either a neuron-like or radial glia-like IPC. Neuron-like cells express and accumulate neuronal factors that will lead to differentiation to two neuronal daughter cells after mitosis. Radial glia-like cells express less total RNA, and fewer neurogenic transcription factors, and most of these cells re-enter the cell cycle. The fate choice is finally executed after mitosis but before again entering the cell cycle.

Next, we examined the maturation axis of cortical development. In the integrated lineage trajectory, different time points were ordered by age, with cells from younger donors residing mostly in the inner part of the cell cycle loops and the upper layers of the neuronal lineage, whereas cells from older donors were found in the outer part of the cell-cycle loop and the bottom layers of the neuronal lineage (Fig. 3H). To quantify this maturation and find associated genes, we performed differential abundance analysis using Milo (*26*). The analysis confirmed the maturation pattern and split the lineage to early and late states (Fig. 3I-J). We tested for differential expression between early and late cells for each class type. To avoid technical batch effects due to 10X chemistry differences, we performed a test using all the samples accounting for batch differences, this test was stringent, hence we also included two differential expression tests that compared early and late states within only Chromium chemistry version 2 samples, and only within 11-14 p.c.w Chromium version 3 samples (fig. S6A-B). These comparisons revealed hundreds of genes that significantly changed in each class type between early and late states (table S6, Fig. 3L, and fig. S6C-D). Interestingly, although most of the genes were differentially expressed in one or two class types (fig. S6C), 28 genes showed higher expression in either early or late state throughout the lineage. These include *FABP7, POU3F2, HTRA1* in late state and *GADD45B, TTR*, DDIT4, and *LEF1* in early state. Previous studies showed that the transcription factors *POU3F2* and *POU3F3* are expressed in the proliferative zones and upper layer neurons in mouse and primate and that they promote the production of the *SATB2* upper layer cortical neurons, further supporting our findings (*27, 28*). This may indicate that this specific set of genes regulates temporal change in radial glia progenitors and instructs the generation of specific types of daughter neurons at specific timepoints.

The greatest number of differentially expressed genes was detected between early and late radial glia and glioblasts (728 genes). Most of those genes showed increased expression level at the late stage. Examination of the genes associated with late radial glia revealed enrichment of genes related to extracellular matrix, focal adhesion, and axon guidance (table S7, fig. S6E). This major transcriptional transformation was previously observed in developing mouse and was predicted to drive these progenitors from internally directed to a more exteroceptive state and also towards gliogenesis (*10, 29*). We compared the mouse and human differentially expressed genes in radial glia between early and late states (*10*), revealing a significant overlap between them, especially between the late genes of both species (Fig. 3K). This observation supports and extends a study by Telley et al. that revealed a conservation of 100 early-late radial glia genes between mouse and human progenitors (*29*). We noted that some of these conserved genes were previously defined as basal radial glia markers including *TNC, HOPX* and *FAM107A* (*30*). Since basal radial glia are rare in mouse (*31–33*), our findings may suggest that those marker genes are part of a broader late radial glia/ glioblasts related program and not specific to basal radial glia. Thus, our results show a conserved program that drive progenitors towards expressing neurogenic and migration-related genes as well as towards gliogenesis.

## Development and migration of forebrain GABAergic neurons

The development of inhibitory interneurons in the ventral telencephalon is largely driven by the spatial distribution of the progenitors in three ganglionic eminences (GEs). To examine the GABAergic inhibitory interneuron lineage, we extracted telencephalic clusters that expressed *DLX2* together with telencephalic clusters that were not defined as excitatory (*FOXG1*+, *EMX1*-) in order to include early radial glia cells that give rise to interneurons excluding OPCs (*PDGFRA*+, *OLIG1*+) (*34–36*). We analysed the dataset for each post conceptional week separately. This resulted in branched trajectories, originating from radial glia/glioblasts and two types of IPCs that branched into lineages clearly defined by ganglionic eminence marker genes and neuronal genes (Fig. 4A, fig S7A-G). We could not detect the neuroblast marker *NHLH1* in these trajectories (Fig. 4A, fig S7A). This confirms findings in the mouse, where *Nhlh1* labeled a distinct neuroblast state at all levels of the neural tube except the ventral forebrain (*10*).

**Figure 4.**
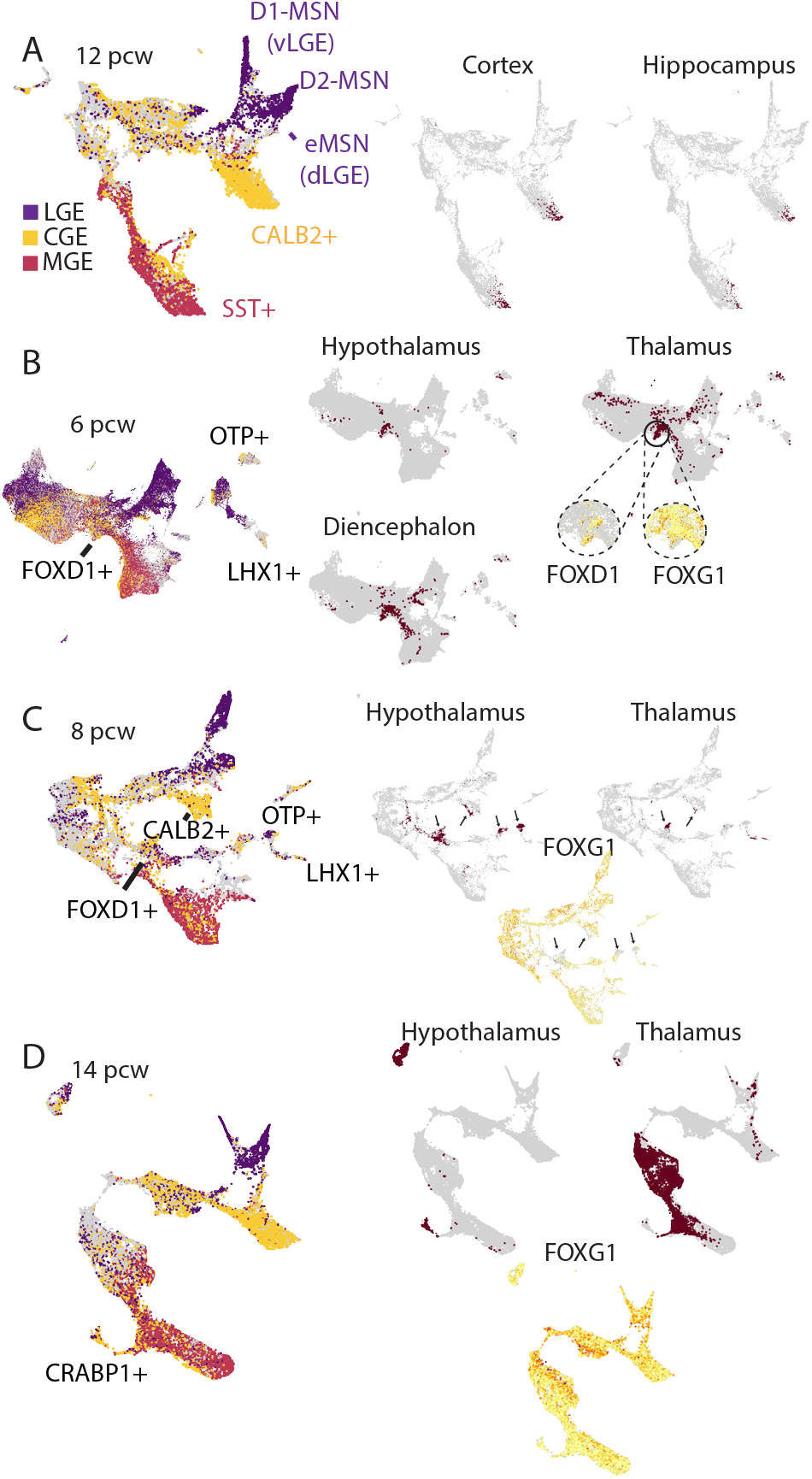
Forebrain GABAergic neurons migration. **(A)** UMAP projections of *DLX2*+ GABAergic lineage cells from 12 p.c.w telencephlic samples (left). Cells are colored according to the expression of different ganglionic eminences marker genes score. Selected interneuron type are marked. Cells from cortical and hippocamapal samples that were clustered with *DLX2*+ cells are colored in brown (right). **(B)** UMAP projections of *DLX2*+ GABAergic lineage cells from 6 p.c.w samples (left). Cells are colored according to the expression of different ganglionic eminences marker genes score. Selected interneuron types are marked. Cells from diencephalon, thalamic and hypothalamic samples that were clustered with *DLX2*+ cells are colored in brown (right). Close-up images show *FOXG1* and *FOXD1* expressing cells. **(C)** UMAP projections of of *DLX2*+ GABAergic lineage from 8 p.c.w samples. Cells are colored according to the expression of different ganglionic eminences marker genes score (left). Selected interneuron type are marked. Cells from thalamic and hypothalamic samples that were clustered with *DLX2*+ lineage cells are colored in brown (upper right). *FOXG1* expressing cells are colored (lower right). Arrows indicate specific locations on the manifold with higher concentration of thalamic or hypothalamic cells and low expression of *FOXG1*. **(D)** UMAP projections of *DLX2*+ GABAergic lineage from 14 p.c.w samples. Cells are colored according to the expression of different ganglionic eminences marker genes score (left). Selected interneuron types are marked. Cells from thalamic and hypothalamic samples that were clustered with *DLX2*+ lineage cells are colored in brown (upper right). *FOXG1* expressing cells are colored in yellow (lower right).

The medial and caudal ganglionic eminences (MGE and CGE, respectively) generate different types of cortical and hippocampal interneurons (*37–41*). Indeed, at the tips of the MGE and CGE trajectories we observed neurons that were dissected from the cerebral cortex and the hippocampus, indicating that they had migrated there (Fig. 4A, fig S7H-I). Cortical MGE-derived neurons were detected at 6 p.c.w and later time points while CGE-derived neurons were observed at 9 p.c.w and later time points. Interneurons and IPCs originating from the lateral ganglionic eminence (LGE) were also observed in cortical dissected samples, however, this was not consistent in all samples and was not reported before, thus suggesting that this resulted from misdissection (the LGE is adjacent to the lateral cortex) (fig S7H-I).

In order to examine the potential migration of telencephalic interneurons to non-telencephalic regions, we broadened the scope by including clusters that expressed *DLX2* from the whole brain together with telencephalic clusters that were not defined as excitatory (*FOXG1*+, *EMX1*-) excluding OPCs (*PDGFRA*+, *OLIG1*+). This yielded similar trajectories but now included cells from non-telencephalic regions. Many of these cells did not express *DLX2* suggesting that they were similar to telencephalic interneurons but probably did not originate in the telencephalon. The vast majority of the *DLX2*-expressing cells were dissected from the diencephalon, thalamus and hypothalamus regions. At 6 p.c.w we identified a subset of *DLX2*- and *FOXD1*-expressing cells in the thalamus (Fig. 4B, fig S8A-B). These cells were clustered with telencephalic radial glia, however they did not express *FOXG1*, indicating that they were transcriptionally similar to GE-derived cells but originated from a non-telencephalic region. Mouse *Foxd1* expression is restricted to prethalamus and hypothalamus during neurogenesis and was recently shown to mark progenitors that give rise to a minor population of *Sox14*-*Pvalb*+ thalamic interneurons (*42–44*). Consistent with this, we detected more than hundred diencephalic and thalamic cells that were clustered with MGE lineage cells and did not express *FOXG1*. These cells did not express high levels of *FOXD1* but they might have originated from *FOXD1*+ progenitors (Fig. 4B, fig S8C). Thus, our findings indicate a similar *FOXD1*+ population of progenitors in human thalamus. At 8 p.c.w we identified *CALB2*-expressing interneurons in the thalamus and hypothalamus, which clustered with CGE-derived neurons (Fig. 4B, fig S8D,E). However, again most of these cells did not express *FOXG1* and therefore probably are not generated in the telencephalon.

At 14 p.c.w. the thalamus sample contained many cells that expressed MGE marker genes and *FOXG1*. The cells were classified as proliferating cells (radial glia, glioblasts, and IPCs) and neurons. The hypothalamus sample contained *DLX2*+ cells that expressed *SST* and *SPARCL1*. The thalamic neurons were clustered in a minor trajectory together with telencephalic neurons expressing *CRABP1* and were not clustered with cortical SST+ neurons (Fig. 4D, fig S8F-G). We had only dissected a single thalamic sample at this stage, hence we cannot rule out that they were caused by misdissection. However, our results are consistent with the report of neuronal migration from the MGE to the thalamus between 15 and 26 weeks of gestation in human and further suggest that these cells give rise to *CRABP1*+ interneurons in the thalamus (*5*).

## Region-specific glioblasts and pre-astrocytes

Astrocytes are heterogeneous along multiple different axes (*45, 46*). The earliest described distinctions are those between fibrous and protoplasmic astrocytes (*47*), and the Müller glia of the retina and Bergmann glia of the cerebellum. More recently, layer-specific cortical astrocytes have been defined using spatial gene expression analysis (*48*). Single-cell analysis has revealed region-specific astrocyte types in the mouse, including two major types residing in the telencephalon and non-telencephalic regions, respectively (*49*), and finer distinctions e.g. between cortex and hippocampus (*50*).

Less is known about the developmental origin of adult astrocyte types. In the developing mouse nervous system, a transitional cell type called glioblast bridges radial glia with nascent astrocytes and oligodendrocyte precursor cells (OPCs). To identify the corresponding human cells, we examined the dendrogram of cell types (Fig. 1) and the distribution of glial cells on two-dimensional embeddings, and searched for late radial glia-like cells oriented away from neurogenesis. Searching for common markers, we found that cells expressing both *BCAN* (*51*) and *TNC* identified putative glioblasts in all brain regions (Fig 1D). Nonetheless, the transition from radial glia to glioblast appeared gradual rather than sudden (Fig. 3), and the distinction therefore somewhat arbitrary.

We found 44 such clusters, which we re-embedded together with oligodendrocyte lineage cells to reveal their transcriptional relationships (Fig. 5 and fig. S9). Most glioblasts were post-mitotic (fig. S9C). They were observed from post-conception week 6, and their heterogeneity was dominated by spatial rather than temporal differences (Fig. 5A and fig. S9b). Minor glioblast clusters were identified as choroid plexus (*FOLR1, TTR*; fig. S9E) and ependymal cells (*PIFO, CCNO*; fig. S9E).

**Figure 5.**
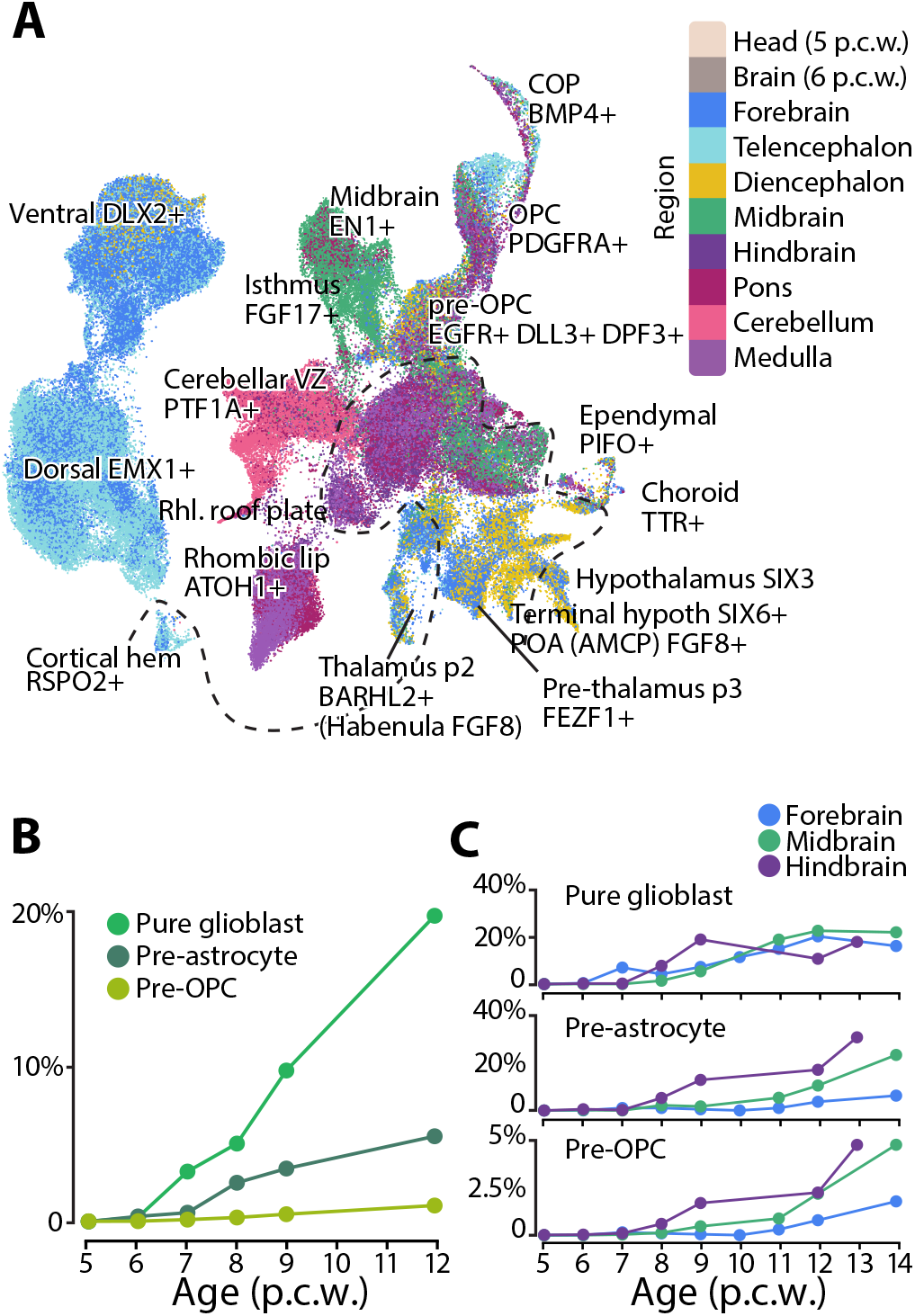
Glioblasts. **(A)** tSNE showing glioblasts, OPCs and COPs, colored by region as indicated. Labels indicate putative spatial position of clusters given the marker genes indicated. Dashed line separate pure glioblasts (above) from pre-astrocytes (below). **(B)** Fraction of pure glioblasts, pre-astrocytes and pre-OPCs detected by age (corrected for sampling). **(C)** Fraction of pure glioblasts, preastrocytes and pre-OPCs detected by age and region (corrected for sampling).

About half of all glioblasts additionally expressed the astrocyte-specific water channel *AQP4* and tight-junction Connexin-43 (encoded by *GJA1*), and we designated those as pre-astrocytes (fig. S9D). Pre-astrocytes were found in the diencephalon, midbrain and hindbrain, and retained region-specific transcriptional identity. Telencephalic pre-astrocytes likely develop after the latest timepoint sampled here, as suggested by low-level *AQP4* expression in late glioblasts from that region (fig. S9D).

One cluster (#510, fig. S9A) was identified as pre-OPCs based on expression of *EGFR* and *DLL3* (*51*), and comprised cells from all major brain regions. Compared with pre-astrocytes and pure glioblasts, pre-OPCs emerged significantly later and in smaller numbers (fig. 5B). All types of glioblasts emerged earlier in hindbrain, followed by midbrain and forebrain (fig. 5C).

Thus the developing human brain comprises a diverse set of glioblasts, differentiating into region-specific pre-astrocytes and OPCs, providing the basis for generating a wide variety of adult macroglia supporting region-specific functions. It seems plausible that region-specific astrocytes reside in nearly every brain region, and that they are developmentally specified to perform distinct region-specific functions in the adult.

## Oligodendrocyte precursors cells defined by their developmental tissue origin

Oligodendrocyte precursor cells (OPCs) are the progenitors to oligodendrocytes, the myelin-producing glial cells that insulate axons and mediate saltatory conduction of the electrochemical impulses between neurons. Although oligodendrocytes have been described to be morphologically (*52*), regionally (*53–55*) and transcriptionally (*56, 57*) diverse, heterogeneity among developing OPCs has not been consistently observed. We previously observed weak evidence of the expression of region-specific patterning transcription factors in adult mouse OPCs, but this was not associated with downstream effector genes and we could not rule out low-level contamination from adjacent neurons (*49*).

Here, we found that human developing OPCs are indeed regionally diverse and exhibit distinct anteroposterior transcriptional identities (Fig. 6A, 6C, 6D, 6E). We identified 8 clusters of the OPC population, a total of 6190 cells, including two clusters that we identified as committed oligodendrocyte precursor cells (COPs) (Fig. 6A). In the hindbrain, OPCs started emerging at 6 p.c.w. followed by a gradual increase in the midbrain and forebrain around 8 p.c.w., consistent with — and even two weeks earlier than — what was recently found in the forebrain by van Bruggen and colleagues (*51, 58*). In the hindbrain OPCs may even emerge slightly earlier, indicated by two cells found in a single 5.5 p.c.w. embryo (Fig. 6B). We also identified committed oligodendrocyte precursor cells (COPs) predominantly in the hindbrain, appearing as early as 6 p.c.w. (Fig. 6B), an even earlier sign of maturation along the oligodendrocyte lineage than what has previously been reported in humans (*59–61*). However, despite the prolonged period of development of OPCs and COPs ranging from 5.5 to 14 p.c.w., we did not observe any mature oligodendrocytes. This suggests that COPs are arrested in their lineage progression, and may serve a function other than myelination during this critical developmental period.

**Figure 6.**
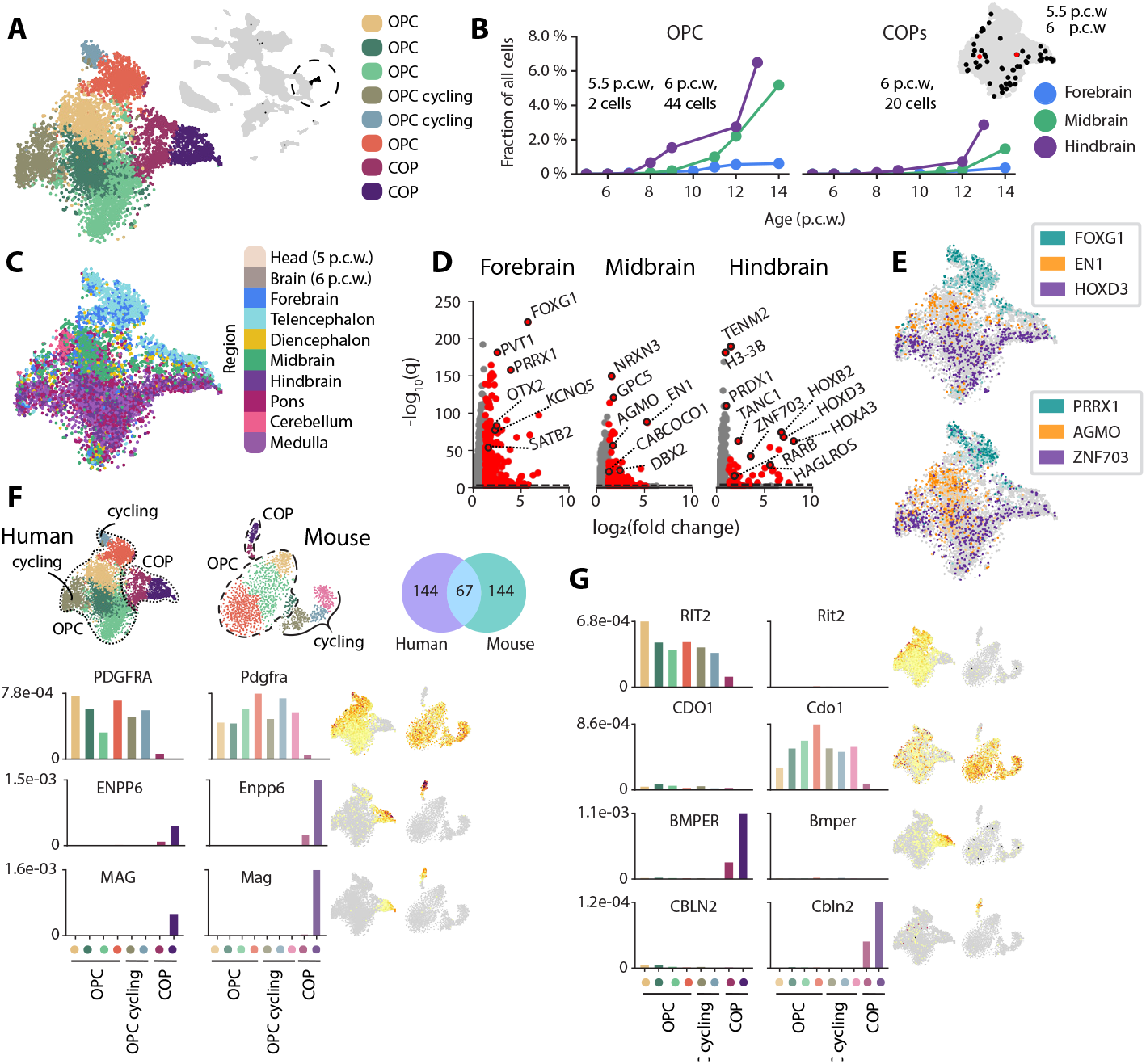
Regional heterogeneity in oligodendrocyte precursors. **(A)** tSNE embedding of all OPCs and COPs, colored by cluster. Small tSNE (grey) showing OPC population in the whole dataset. **(B)** Distribution of ages within OPC and COP groups, showing fraction of cells per age and main region (forebrain, midbrain, hindbrain). Small tSNE (right) displaying number of cells from 5.5 p.c.w (red) and 6 p.c.w (black), zero expression shown in grey. **(C)** tSNE of all OPCs and COPs colored by region. **(D)** Top differentially expressed genes in fore-, mid- and hindbrain within the OPC population. Genes selected with a *q*-value < 10e-40 (dashed line). **(E)** Example of top differentially expressed genes, marking each region, top: *FOXG1* (forebrain), *EN1* (midbrain) and *HOXD3* (hindbrain). Left: *PRRX1* (forebrain), *AGMO* (midbrain), *ZNF703* (hindbrain). **(F)** Comparison between OPCs and COPs in human and mouse respectively. Individual clusters show OPC, COP and cycling cell populations in both species. Venn diagram of differentially expressed genes (significance level) between OPC and COP cells within human and mouse, with 67 genes shared between the species (lightblue). Commonly used gene markers for OPCs (*PDGFRA*), COPs (*ENPP6*) and more maturing COPs (*MAG*) were part of the shared list of differentially expressed genes and expressed in both species (bar plots and miniature tSNEs). **(G)** Differentially expressed genes that are specific to either human: *RIT2* (OPCs) and *BMPER* (COPs) or mouse: *Cdo1* (OPCs) and *Cbln2* (COPs), illustrated in bar plots. Miniature tSNEs showing expression patterns of the same genes.

To further understand the regionalization of the OPCs and COPs, we asked if their diversity could be explained by any classical patterning genes typically expressed in the fore-mid- and hindbrain (Fig. 6D, S10C). We performed differential gene expression analysis between cells from each region within the OPCs and COPs separately (see differential test groups Fig. S10C). This resulted in a list of differentially expressed genes (see table S8) per region, some of them being developmental regional patterning genes such as *FOXG1* (forebrain), *EN1* (midbrain-hindbrain boundary) and *HOXD3* (hindbrain) (Fig. 6E, fig. S10C, S10D). Beyond the patterning genes (fig. S10D) a major fraction of the region-specific genes were transcription factors, ion channels and receptor subunits (Fig. 6D, 6E, S10C, table S8). Although both clusters of cycling OPCs were dominated by cell cycle genes, they retained a transcriptional profile that separated them into forebrain- and hindbrain-specific cycling OPCs (fig. S10C). This finding shows that there are multiple sources of proliferating OPCs that give rise to subsequent maturing OPCs, demonstrating that their regional identity is already established before they become post-mitotic.

The expression of *PRRX1, AGMO* and *ZNF703* illustrates the regionality observed in the OPC group (Fig. 6E, bottom). *PRRX1*, a transcription factor typically expressed in mesoderm-associated tissues and that is involved in craniofacial and limb development, was specific to the forebrain-derived OPCs (Fig. 6E). In addition, part of the forebrain COP trajectory showed expression of *PRRX1* (Fig. 6E). A recent study described its role in regulating cell-cycle progression by inducing (reversible) quiescence in cultured human fetal primary OPCs (*62*). These cells underwent reduced proliferation and migration upon *PRRX1* expression as opposed to differentiated OPCs that showed a downregulation of the same gene. In our data, *PRRX1* was mostly expressed in forebrain OPCs including the cycling OPCs of this region (Fig. 6A, 6E, S10B), suggesting a similar involvement of *PRRX1* in the regulation of OPC differentiation as previously described (*62*). *AGMO* is an enzyme that catalyzes the cleavage of ether bonds in ether lipids such as alkylglycerols, while *ZNF703* is a zinc-finger protein that acts as a transcriptional repressor and has been shown to be important for hindbrain formation and generation of neural crest progenitors in the *Xenopus* embryo (*63, 64*). Together, these genes (along with the other regionally differentially expressed genes) illuminate their potential implication in neurodevelopmental disorders.

## Conserved and divergent properties of OPCs

After noticing the heterogeneity of OPCs, we sought to understand if this transcriptional pattern was conserved between species. Comparisons between species are challenging because of difficulties in aligning their anatomical, functional and temporal differences. For example, while mice have a gestation period of roughly three weeks with an onset of myelination postnatally, the same process is thought to occur during the last trimester in humans (*65*). Despite differences in sampling depth, experimental methods used, and clustering algorithms, we were able to match OPCs, cycling OPCs and COPs to the corresponding cell states in our previously published developmental mouse dataset (*10*) (Fig. 6F, fig. S10E). In the mouse, these cells included timepoints from E12 to E18 (until birth) which in humans roughly correspond to 5 p.c.w. up to at least 8 p.c.w. and onwards. We assumed that we covered overlapping developmental ages of the species and considered it comparable. To enrich for functionally relevant genes, we first identified those genes differentially expressed between OPCs and COPs in each species, and then compared the resulting gene sets between the species (Fig. 6E, 6F, fig. S10E, table S9). 67 of 211 (32%) differentially expressed genes were shared between species (Fig. 6F, fig. S10F). Many prototypical marker genes known in the early oligodendrocyte trajectory were conserved, such as *PDGFRA* (OPCs), and *ENPP6* and *MAG* (COPs; Fig. 6E, fig. S10F). The remaining 144 genes in each species were not shared, and their expression patterns either diverged or the gene was not expressed in the other species (fig. S10F). For example, *RIT2* (OPCs) and BMP-binding endothelial regulator (*BMPER;* COPs) clearly distinguished the two populations in human, as opposed to the mouse where *Cdo1* and *Cbln2* labeled the same groups (Fig. 6G, fig. S10F). *BMP4* was enriched in COPs in both species, but more highly expressed in mouse COPs (fig. S10F). *BMPER* is involved in BMP-signaling and acts as an inhibitor for BMP proteins (like BMP2 and BMP4; ref). Studies in other model organisms have shown that the inhibition of BMP-signaling is necessary for the commitment of the oligodendrocyte lineage and that overexpression of BMPs causes a reduction in the number of oligodendrocytes (*66–68*). Hence, the expression of *BMPER* observed in the human but not mouse OPCs demonstrates a species-specific mode of BMP-signaling, which may regulate the rate of differentiation of OPCs. Altogether, while we observed conservation of many functionally relevant genes between mouse and human in the oligodendrocyte lineage, the majority of differentially expressed genes showed a species-specific expression pattern (Fig. 6F, 6G, fig. S10).

## Discussion

We have presented a comprehensive transcriptomic survey of human brain development during the first trimester. There are several important caveats. For example, due to the nature of human samples, it was impossible to obtain timed samples, resulting in uneven coverage of temporal processes. Often, not every region in a specimen was free of damage, leading to missing regions which could only be partially compensated for by other specimens of similar age. We used freshly dissociated cells, but the length of time before sample collection, and the ambient temperature and buffer conditions during that time could only be partially controlled. Inordertominimizethedelaybetweensample collection and cell capture, we used coarse dissections, which may sometimes not have coincided perfectly with neuroanatomical boundaries. Nevertheless, we provide here a first comprehensive cell census of early human brain development. The results expand our knowledge especially of glial cell development, and underscores the fundamental similarity between all major cell classes emanating from the neural tube: neurons, astrocytes and oligodendrocytes. All three classes develop from region-specific radial glia, and transition through intermediate cell states — neuroblasts, glioblasts, neuronal IPCs, pre-astrocytes and pre-OPCs — that retain all or most of that patterning. In a separate paper (Siletti et al. *submitted*) we demonstrate very similar patterns of region-specific glia in the adult human brain. It seems that the human brain makes use of highly regionally and locally specialized neurons, supported by similarly region-specific astrocytes and oligodendrocytes. The functional implications of this view remain to be uncovered, but it is worth highlighting that many neuronal disorders implicating glia show strong region-specific patterns of occurrence. For example, pediatric high grade gliomas show distinct mutations associated with occurrence at specific neuroanatomical sites (*69*). For example, H3.1 K27M mutations are prevalent in the thalamus, pons and spinal cord, whereas H3.3 G34V/R mutations occur more predominantly in tumors located to the cortex and striatum. It will be interesting to see if region-specific glial cell types can explain some of these phenomena. More generally, our cell census provides a rich resource for exploring normal human brain development, and a powerful reference for interpreting diseased brain tissue.

## Acknowledgments

We gratefully acknowledge the services of the Developmental Tissue Bank (Department of Neurobiology, Care Sciences and Society, Karolinska Institutet) core facility in providing prenatal tissue. We acknowledge support from the National Genomics Infrastructure in Stockholm funded by Science for Life Laboratory, the Knut and Alice Wallenberg Foundation and the Swedish Research Council, and SNIC/Uppsala Multidisciplinary Center for Advanced Computational Science for assistance with massively parallel sequencing and access to the UPPMAX computational infrastructure. We thank Kimberly Siletti (Karolinska Institutet), Amit Zeisel (Technion, Israel), Eneritz Agirre, Petra Kukanja (Karolinska Institutet) and David van Bruggen (Hardvard University) for critical input and helpful discussions.

## Author contributions

SL conceived the project. EB and LH performed single-cell experiments and sequencing libraries. LH processed libraries for sequencing. PL handled data acquisition from sequencing. LEB performed the EEL FISH experiment, analyzed and visualized the data. EV performed anatomical annotation of the spatial EEL data. KWL and EA annotated midbrain clusters. XL and ES dissected and supplied fetal tissue. XH and RB dissected and supplied fetal tissue. ZA performed Visium experiments with supervision by JL. SL, MDG and EB analyzed, visualized the data and wrote the manuscript.

## Data and materials availabilit

Raw sequencing data is available from the European Genome Phenome Archive (accession EGAS00001004107). Jupyter notebooks for reproducing figure panels, and links to the annotated expression count matrix are available at https://github.com/linnarsson-lab/developing-human-brain/.

## Methods

### Human tissue collection

For tissue collected at Karolinska Institute, patients seeking abortion at the gynecology clinic were asked about their interest in donating the aborted tissue to research. Patients that agreed, signed a written consent after receiving information, both written and oral, given by a physician or midwife. The use of abortion material was approved by the Swedish Ethical Review Authority and the National Board of Health and Welfare. Age (post-conception) of the embryos and fetuses was estimated using clinical information (last menstrual period, ultrasound), true crown-rump-length and anatomical landmarks (*70, 71*). After the abortion the tissue was immediately transported to the laboratory and dissected in ice-cold 0.9% NaCl solution. Most of the calvarium was cut away, and the brain was carefully lifted out of the skull base after separating the medulla oblongata from the cervical spinal cord. For scRNAseq, different CNS regions were dissected and kept in ice-cold Gibco Hibernate-E medium (ThermoFisher) until further processing. For spatial analysis brains were covered by Tissue-Tek O.C.T. Compound (Sakura Finetek) in cryomolds, snap-frozen in a slurry of 2-methylbutane (Sigma-Aldrich) and dry ice, and stored at -80°C pending sectioning.

For tissue collected in Cambridge, donated foetal tissue stored in Hibernate-E medium (Hib-E, Thermo Fisher Scientific A1247601) was collected from local maternity hospital soon after passing. The relevant parts of brain tissue were dissected in a class II hood on the day of collection and stored in Hib-E overnight in a fridge. The tissue was shipped to Sweden at refrigerated temperature the next day, normally delivered two days after abortion. The procedure is covered under ethics REC: 96/085. Subsequent processing was done as described for Karolinska samples.

### Cell dissociation

Brain tissue was processed around 6 to 48 hours post tissue collection, depending on source (Cambridge/ Karolinska Hospital). Tissues that were not processed within the same day of collection were stored at 4°C in Hibernate E medium (ThermoFisher) during transport or overnight. A carbogenated (95% O_2_ / 5% CO_2_) ice cold Earle’s Balanced Salt Solution (EBSS) was used throughout the whole procedure. All brain regions were dissociated separately using the Worthington’s Papain Dissociation System (Worthington) (Protocols. io, https://dx.doi.org/10.17504/protocols.io.xmbfk2n). Tissues were enzymatically digested at 37 °C for 10 to 30 min (depending on the developmental time point, younger ages kept shorter), followed by regular trituration using fire polished glass Pasteur pipettes. Cell suspensions were filtered through a 30 μm cell strainer (CellTrics, Sysmex), centrifuged for 5 min at 200 g to obtain cell pellets, followed by careful removal of the supernatants and resuspension in EBSS (Worthington), using as small volume as possible depending on tissue size and cell density. Cell concentrations were estimated using a counting hemocytometer (Bürker/Neubauer chamber) and diluted with EBSS until the desired concentrations were reached. All suspensions were kept on ice until the next step of loading the cells on the 10X Chromium chips.

### Single-cell RNA sequencing

Single cells were captured using the droplet-based single-cell RNA sequencing platform Chromium (10X Genomics). Roughly half of the sampling was done using the Chromium Single Cell 3’ Reagent Kits Version 2 and the other half with Version 3 (see detailed sampling in table S1). Cell suspensions were adjusted to concentrations between 800 - 1200 cells/μl, targeting 3000 - 5000 cells per reaction. cDNA synthesis was performed with 12 PCR cycles and the rest of the library preparation was performed according to the manufacturer’s instructions (10X Genomics, Illumina). All libraries were sequenced on an Illumina NovaSeq 6000 using S4 to a target sequencing depth of 100,000 reads/cell. Sequencing saturation was examined for each sample using preseq (https://github.com/smithlabcode/preseq). Any samples that were not saturated to 60% were sequenced more deeply as needed, using preseq predictions.

### Enhanced Electric fluorescent in-situ hybridization (EEL)

440 genes were selected to cover major cell types in the human developmental brain using single-cell RNA-seq data and literature (table S3). EEL FISH was performed as previously described (*8*) with a minor change in the digestion step, that was performed two times 10 minutes with a concentration of 1 U/ml of proteinase K.

### Standard Visium Spatial Gene Expression library preparation

Fresh-frozen human embryo sample was cryo-sectioned at 10 μm thickness. Sections were placed onto 10X Genomics Visium arrays and stored in -80°C before processing. Spatial gene expression libraries were prepared following 10X Genomic Visium Gene Expression protocol (https://assets.ctfassets.net/an68im79xiti/2q34xwfHy2nbeFlH47BlOq/ffa48b53627a582c6b7f2c9fd90af91e/CG000239_Visium_Spatial_Gene_Expression_User_Guide_Rev_F.pdf). Finished libraries were sequenced on Illumina Nextseq2000 according to manufacturer’s instructions (read 1, 28 bp; read 2, 90 bp).

## Data preprocessing

Sequencing runs were demultiplexed with cellranger mkfastq version 4.0.0 (10x Genomics) and filtered through the index-hopping-filter tool version 1.1.0 (10x Genomics). Unique molecular identifier (UMI) counts were determined using STARSolo version 2.7.10a with the following parameters:

- -soloFeatures Gene Velocyto
- -soloBarcodeReadLength 0
- -soloType CB_UMI_Simple
- -soloCellFilter EmptyDrops_CR %s 0.99 10 45000 90000 500 0.01 20000 0.01 10000
- -soloCBmatchWLtype 1MM_multi_Nbase_pseudocounts
- -soloUMIfiltering MultiGeneUMI_CR
- -soloUMIdedup 1MM_CR
- -clipAdapterType CellRanger4
- -outFilterScoreMin 30

Barcode whitelists were downloaded from the 10x Genomics website. Exonic, intronic, and ambiguous counts were summed for clustering analysis.

The reference genome and transcript annotations were based on the human GRCh38.p13 gencode V35 primary sequence assembly. However, we filtered the reference. In this study, only reads that were uniquely aligned to one gene were counted. Thus, all the reads that aligned to more than one gene were lost and the related genes had lower or zero counts.

To lower the rate of this read loss, we used the human GRCh38.p13 gencode V35 primary sequence assembly from which we discarded genes or transcripts that overlapped or mapped to other genes or non-coding RNAs 3’ UTR, leaving only one of these transcripts in the genomic reference. We used BLAST to align the last 400 nt (3’ UTR) of all protein coding transcripts and non-coding transcripts to all other genes (maximum 4 mismatches, minimum alignment length 300 nt). We resolved all the matches by the following procedure:

1. Fusion genes - fusion genes were filtered based on their names: genes with names that contained both fusion genes [‘gene1-gene2’] were discarded.
2. Non-coding transcripts - non-coding transcripts that matched another coding transcript were discarded.
3. Overlapping transcripts (both examined transcripts were either coding or non-coding)

a. If one of the examined genes name matched the pattern ‘XX######.#’ its transcript was discarded.
b. If one of the examined transcripts belonged to a gene where one or more of its transcripts already got discarded during the procedure, it was discarded as well.
c. We discarded transcripts of the gene with a lower number of splice variants.
d. Otherwise, the transcript that its 3’ UTR overlapped the other transcript was kept in the genomic reference.
4. Paralogs - we mapped all related genes (that aligned to one another) and selected one of the paralogs with the highest number of splice variants. All other highly similar paralogs were discarded. In special cases, we manually chose the gene.

Altogether, this yielded a new assembly in which we filtered 387 fusion genes, 1140 overlapping transcripts, 414 non-coding transcripts, 1127 coding paralogs, and 350 non-coding paralogs.

### Quality control / preprocessing

Initial quality control of each 10X Chromium sample as a whole was performed by manually examining the result of library preparation (deciding whether to sequence the sample at all) and the output of the primary analysis pipeline (deciding if the sample as a whole was of sufficient quality to be included at all). Samples that failed at these early stages are listed in Table S1 but no cells from them are included in the main dataset.

The result of a Chromium experiment is a raw expression matrix over genes and droplets. For a high-quality sample, most droplets will have contained a single viable cell. However, some droplets may have contained a doublet or a piece of debris. In order to distinguish these possibilities and identify high-quality single-cell droplets, we developed the following algorithm. It is similar in spirit to DropletQC (*72*), but was developed independently by us. First, we used the DoubletFinder (*73*) algorithm to mark putative doublets. Next, we used the relationship between total UMI count and the fraction of unspliced UMIs (fig. S2A-D) to classify droplets as cells, large cells (potentially multiplets), cytoplasmic debris (low total UMI, low unspliced fraction), cellular debris (low total UMI, normal unspliced fraction), nuclear debris (low total UMI, high unspliced fraction) and mitochondrial debris (high fraction mitochondrial reads). To classify droplets, we first selected droplets likely to be single cells by including only droplets with unspliced fraction greater than 0.1, total UMIs greater than 1500 and log(total UMIs) greater than log(200) + log(1000) * unspliced fraction. Next, we fit a two-dimensional gaussian maximum likelihood estimate to the selected droplets (using the logarithm of the total UMIs) and then calculated the probability density function for all droplets, retaining only those with probability greater than 0.1 (except those flagged as doublets). The remaining droplets were then classified as illustrated in fig. S2B based on their location relative to the good cells.

### Cytograph/Cytograph-shoji pipeline

We used the latest version of our in-house cytograph pipeline, which implements standard analysis steps in a modular and scalable fashion. The version used here is called cytograph-shoji and uses a custom tensor database called shoji. However, the analysis steps are standard and can also be performed using publically available single-cell analysis tools such as Scanpy and Seurat.

For the initial clustering, we pooled all cells that had passed quality control. We selected 1000 most variable genes using support-vector regression on the CV-vs.-mean relationship, excluding immediate-early genes, cell cycle genes, mitochondrial genes, and non-coding RNA (the exact lists of genes excluded are available in the cytograph-shoji code, file “species/human.py”). We performed principal component analysis and retained up to 50 components, but only as many as would explain 50% of the variance. We then transformed the PCA using Harmony (*7*) to correct for batch effects due to the 10X Chromium chemistry version (v2 or v3). We computed the manifold as a balanced *k*-nearest neighbors graph with *k* = 25 and euclidean distance metric. We performed initial clustering on the manifold using the Leiden (*74*) algorithm, then removed clusters with less than 25 cells. We trained a support-vector classifier (using stochastic gradient descent with hinge loss) on the remaining cluster labels, and then applied the classifier to the orphan cells to assign their cluster identity to one of the retained (not too small) clusters. We also computed the classifier probability for each cell, as well as the second-best cluster label, which could be used to judge the quality of clusters and transition zones between adjacent clusters.

We then computed a dendrogram of the initial clusters and cut this to yield 40 subsets. Each subset was reclustered using the same procedure as above, and then all the resulting clusters were pooled again to make the complete dataset corresponding to table S2. A few clusters were manually identified as doublets based on inconsistent marker expression, high average doublet score and/or high average UMI count.

We computed gene enrichment, trinarization scores, auto-annotation and cell cycle scores as previously described (*10*). We used the expression of a set of well-known cell cycle genes (*75*) as a proxy for active proliferation. We calculated the cell cycle score as the fraction of UMIs those genes represented, and used a threshold of 0.4% to call a cell cycling (fig. 3A-C). Similarly, a score for each phase of the cell cycle was calculated for each cell using a subset of known genes expressed in that phase.

**Gene enrichment** is a measure of overexpression in a cluster relative to other clusters, taking into account both mean expression and fraction of non-zero cells.

**Trinarization** is a measure of the probability *P* that a gene is expressed in more than 20% of the cells, resulting in calls of ‘not expressed’ (*P* < 0.05), ‘ambiguous’ (0.05 ≤ *P* ≤ 0.95) or ‘expressed’ (*P* > 0.95). To determine if a gene, or a set of genes, was expressed in a cluster, we used the product of the trinarization score (i.e. the joint probability) with a cutoff of 0.95. Generally, when we say that a gene is expressed in a cluster, this is the formal method we used to support that statement.

**Auto-annotations** are well-known sets of gene markers applied automatically during the analysis pipeline, using the same trinarization method.

**Cell classes** were defined using auto-annotation with the markers described in the main text. When a cluster occasionally received multiple class labels, they were resolved by priority: Fibroblast > Immune > OPC > Glioblast > Radial glia > Neuroblast > Neuron > (other). When a cluster received no class label, one was assigned manually where possible. Neuronal IPCs were defined as proliferating Neurons and Neuroblasts.

The excitatory lineage analysis in Fig. 3 was performed before the cytograph-shoji pipeline was completed, and therefore used a previous version of cytograph (*10*). In brief, we used 10X Genomics cellranger pipeline (version 3.0.2) to align the reads from all telencephalic samples. RNA counts were attributed to spliced and unspliced transcripts by running velocyto (*76*) (version 0.17.11) with standard parameters. For initial classification we used PCA and transformed it using Harmony to correct for batch effect resulting from 10X Chromium chemistry versions. A radius nearest neighbour graph (RNN) was then computed using the information radius (also known as the Jensen–Shannon divergence (JSD)) to link cells with near-identical gene-expression states. This RNN graph was clustered using Louvain algorithm. For cortical excitatory lineage analysis, we selected clusters that expressed *EMX1*.

### Spatial mapping of single cell data

The anatomical atlas was made in Napari (*77*). Mean expression profiles of single cell RNA-seq clusters of cells from 5 week old samples were manually selected and mapped to the hexagonally binned spatial data using FISHscale and Bonefight (*8, 10*). Mappings were locally smoothened using a distance weighted average, and the clusters with the highest probability for each spatial hexagonal bin were called to make the cluster location map.

Visium libraries were processed using Space Ranger software from 10X Genomics (version 1.3.1). Reads were aligned to the pre-built human reference genome (GRCh38).

### Cortical excitatory lineage analysis

#### Integration

For this analysis we subset cells from telencephalic samples that were clustered in clusters that expressed EMX1. We used scanpy package for most of this analysis unless mentioned otherwise (*78*). The cortical subregion and the donor ID for each cell was considered as the technical covariates to correct for. We selected 1000 highly variable genes for the Scanorama analysis and 700 highly variable genes for the scVI analysis using scanpy.pp.highly_variable_genes based on ‘seurat_v3’ flavor (*79*) and performed dimensionality reduction and batch correction for each p.c.w. using Scanorama (*14*) and for all time points using the scVI model (*15*) as implemented in scvi-tools (version 0.15.5) (*80*).

To keep a consistent analysis, samples that did not presented batch differences (5,7, and 14 p.c.w) were integrated with the consecutive p.c.w samples (except 14 p.c.w. samples that were integrated with 13 p.c.w samples). We then computed nearest neighbors, clustering using Louvain algorithm and UMAP projection using similarity in the scVI/Scanorama embedding.

#### Cell cycle phase determination

Cells in G1 phase were defined as cells with G1 score > 0.002, cells in S phase were defined as cells with S score > 0.002, and cells in G2/M phase were defined as cells with G2M score > 0.03. In case of ambiguous labels per cell, we resolved it by priority: G2M > S > G1. All other radial glia and IPC progenitors were labeled as non-cycling.

#### RNA velocity and DeepCycle analysis

To compute RNA velocity we used scvelo package (version 0.2.4) (*81*) and run stochastic mode analysis on each batch (cortical subregion or donor as needed) that had more than 1000 cells in every p.c.w. scvelo velocity length was plotted using Scanorama inferred UMAPs for each p.c.w. For fig. S5E we chose one representative batch for each p.c.w. To infer high resolution cell cycle trajectories DeepCycle was used based on scvelo run for subset of IPCs cells for each batch. IPCs were defined based on score calculate by scanpy.tl.score.genes given to each cell by the expression of EOMES, NHLH1 and NEUROD4, thus also including early differentiating IPCs. We could not infer a clear cell cycle trajectories for all examined batches. Eventually, we used the trajectory obtained for donor XDD:313 forbrain samples from 8.5 p.c.w and donor XDD:385 14 p.c.w. Cell cycle phases were inferred for trancriptional phase Θ range using the score given for each cell cycle phase (fig. S5F,H bottom). Mitosis was inferred as the transcriptional phase where there was a sharp drop in UMI counts.

#### Differential abundance analysis

Differences in cell abundances associated with post conceptional age was tested using the Milo framework (*26*), with the Python implementation milopy (https://github.com/emdann/milopy). We tested differential abundance in 3 subsets of cells: (A) all cortical excitatory lineage cells (fig. S6A) (B) 11-14 p.c.w cells of v3 Chromium chemistry samples (fig. S6B, top) (C) 6-10 p.c.w. cells of v2 Chromium chemistry samples (fig. S6B, bottom). We constructed a KNN graph using similarity in the scVI embedding (k = 150 for A (v2 and v3 cells), k = 50 for B (v3 test), and k=100 for C (v2 test)). We assigned cells to neighborhoods on the KNN graph using the function milopy.core.make_nhoods (prop = 0.1).We defined early neighborhoods (SpatialFDR < 0.05, logFC < -0.5) and late neighborhoods (SpatialFDR < 0.05, logFC > 0.5) and tested for differential expression between cells from each of these groups (see ‘Differential gene expression analyses’).

### Region classifier

A random forest classifier from sklearn (RandomForestClassifier) was used to predict the “contamination” of cells belonging to other regions than the dissected ones. The classifier was trained on all brain regions except for cells labelled “Brain” or “Head” with each region downsampled to 70 000 cells per region resulting in a training set of 558 702 cells and 1000 (most variable) genes. The whole dataset (1 665 937 cells) was tested on the trained model and by treating the dissected region labels as the true labels, an accuracy score could be estimated to 0.8. Assuming the model performed well enough, we used it to predict regions per cell and the fraction of the most commonly predicted region was calculated for each individual sequencing library. Following hyperparameters were tuned and used in the final training of the classifier: min_samples_leaf = 3, max_features = 50, n_jobs = -1.

### Differential gene expression analyses

All differential gene expression tests were performed using the diffxpy package (https://github.com/theislab/diffxpy). The Wilcoxon rank-sum test was used unless mentioned otherwise.

#### Cortical excitatory lineage neuroblasts vs. neurons

We normalized raw gene read counts by sequencing depth in each cell using scanpy.pp.normalize_total function and performed log(x)+1 transformation. Differential expression test was performed between neuroblasts and neurons. We considered genes as significantly differentially expressed if the test FDR < 1e-30, mean expression > 0.01 and a log2(fold change) > 1.5 or log2(fold change) < -1.5. The results for the differential expression analysis between cortical excitatory neuroblasts and neurons are provided in table S4.

#### Cortical excitatory lineage neuronal-like IPCs vs. radial glia-like IPCs

We normalized raw gene read counts by sequencing depth in each cell using scanpy.pp.normalize_total function. To find genes related to neuronal-like IPCs and radial glia-like IPCs, we first compare radial glia/ glioblasts cells to differentiating IPCs (fig. S5L, top) to detect genes that are distinctive for these cell types. We further used the genes that were differentially expressed (FDR < 1e-30, and a log2(fold change) > 2 or log2(fold change) < -2) to find those that are also differentially expressed between cells entering the IPC cell cycle and differentiating IPCs (fig. S5L, bottom). We considered genes as significantly differentially expressed if the test FDR < 1e-30 and a log2(fold change) > 2 or log2(fold change) < -2. Genes with log2(fold change) > 2 were further used for heatmap plots (fig S5M,N). The results for this differential expression analysis are provided in table S5.

#### Cortical excitatory lineage early vs. late state

After testing for differences in cell abundances associated with post conceptional age (see ‘differential abundance analysis’ section) we defined each cell as early, late, or not differentially abundant (DA) based on its neighborhood definition (fig. 6A,B).

We then performed differential expression analysis between early and late cells of the main cortical excitatory lineage (excluding cells from the minor Cajal Rezsius lineage) (Fig. S6A,B). To overcome the Chromium chemistry versions bias we used 3 differential expression tests and integrated the resulting differential expressed genes as follows:

A. Early vs. late cells in each major class type were compared (fig. S6A). Wald test was performed and we accounted for chemistry in the test model. We considered genes as significantly differentially expressed if their mean expression > 0.01, the test FDR< 0.05 and log2 fold change > 1 or log2 fold change < -1.
B. Early vs. late cells from 11-14 p.c.w Chromium chemistry v3 samples in each major class type were compared (fig S6B, top).
C. Early vs. late cells from Chromium chemistry v2 samples in each major class type were compared (fig S6B, bottom)

For both B and C tests, we normalized raw gene read counts by sequencing depth in each cell using scanpy.pp.normalize_total function.Wilcoxon rank-sum test was used and the gene was considered as significantly differentially expressed if its mean expression > 0.01, the test FDR<1e-15, and a log2 fold change > 1.2 or log2 fold change < -1.2.

To integrate the results we used the following procedure: For each class:

- All the differentially expressed genes from test A were obtained in the final table.
- In case the gene was not significantly differentially expressed in A, and was significantly differentially expressed in B and C in a consistent manner, the gene was assigned as either ‘early’ or ‘late’ based on the type of cells it was upregulated in, otherwise it was assigned as ‘mixed’.
- If the gene was only significantly differentially expressed in B (late samples, v3), it was assgined as differentially expressed (‘late’) only if it had higher expression in late cells. Otherwise, it was assigned as ‘mixed’.
- If the gene was only differentially expressed in B (early samples, v2), it was marked as differentially expressed (early) only if it had higher expression in early cells. Otherwise, it was assigned as ‘mixed’.

If the pattern of the change was inconsistent between different classes, the gene was assigned as ‘mixed’. HLA genes and gender-related genes were further removed from the final set of differentially expressed genes. The summary table for these differential expression analyses results is provided in table S6.

GO term and pathway enrichment was performed using the implementation of the EnrichR workflow (*82*) in the gseapy Python package (https://pypi.org/project/gseapy/). The results for this analysis are provided in table S7.

#### Oligodendrocyte precursor cells

The test was applied on the raw gene expression matrix (without any prior log-normalization) where each cell was scaled to the median total molecules. For the mouse and human comparison, gene expression values were specifically scaled to 4000 molecules per cell (in between the median total molecules per species, human: 6065 UMIs versus mouse: 3824 UMIs). In the species comparison, all cells from the OPCs and COPs were compared within each species. Differentially expressed genes were selected by a cut-off of following parameters: *q-*value < 10-3 (mouse), *q-*value < 10^−149^ (human), mean expression > 0.01 and a log_2_ fold change ≥ 1.5 (absolute value). Different cutoffs were used per species to compensate for differences in number of cells sampled and the sensitivity of the single-cell chemistry used, and were adjusted to select roughly equal number of genes. For the regional comparison of the OPCs and COPs, all cells from each dissected region were tested against the rest of the cells from the other regions (excluding the cycling OPCs). Differentially expressed genes were selected based on these thresholds: *q-*value < 10^−39^, mean expression > 0.01 and a log_2_ fold change > 1.2 (selecting only up-regulated genes). The results of the analyses are provided in table S8 (region-specific OPCs) and table S9 (species comparison of OPCs).

### Morphometric volume normalization

When estimating the proportion of classes of cells (e.g. as in Fig. 1B), normalization was necessary. We did not sample regions and timepoints uniformly, and therefore a raw estimate would be skewed by over- and under-sampling. In addition, the brain as a whole grows in volume, and different regions grow at different rates. In particular, the telencephalon expands dramatically during the period we sampled. We therefore normalized our sampling to the expected tissue volume (excluding ventricles) based on morphometric estimates of forebrain, midbrain and hindbrain size by age (*83, 84*). We omitted those timepoints where we lacked complete coverage (e.g. when hindbrain was sampled only as pons, omitting medulla); this is why 13 and 14 p.c.w. are missing from Fig. 1B.

### Illustrations

The embryo image in Fig. 1A is a rendering of an actual human embryo, overlaid on a 3D volume representing the nervous system, redrawn from data provided by HDBR (*85*).

**Supplementary Figure 1.**
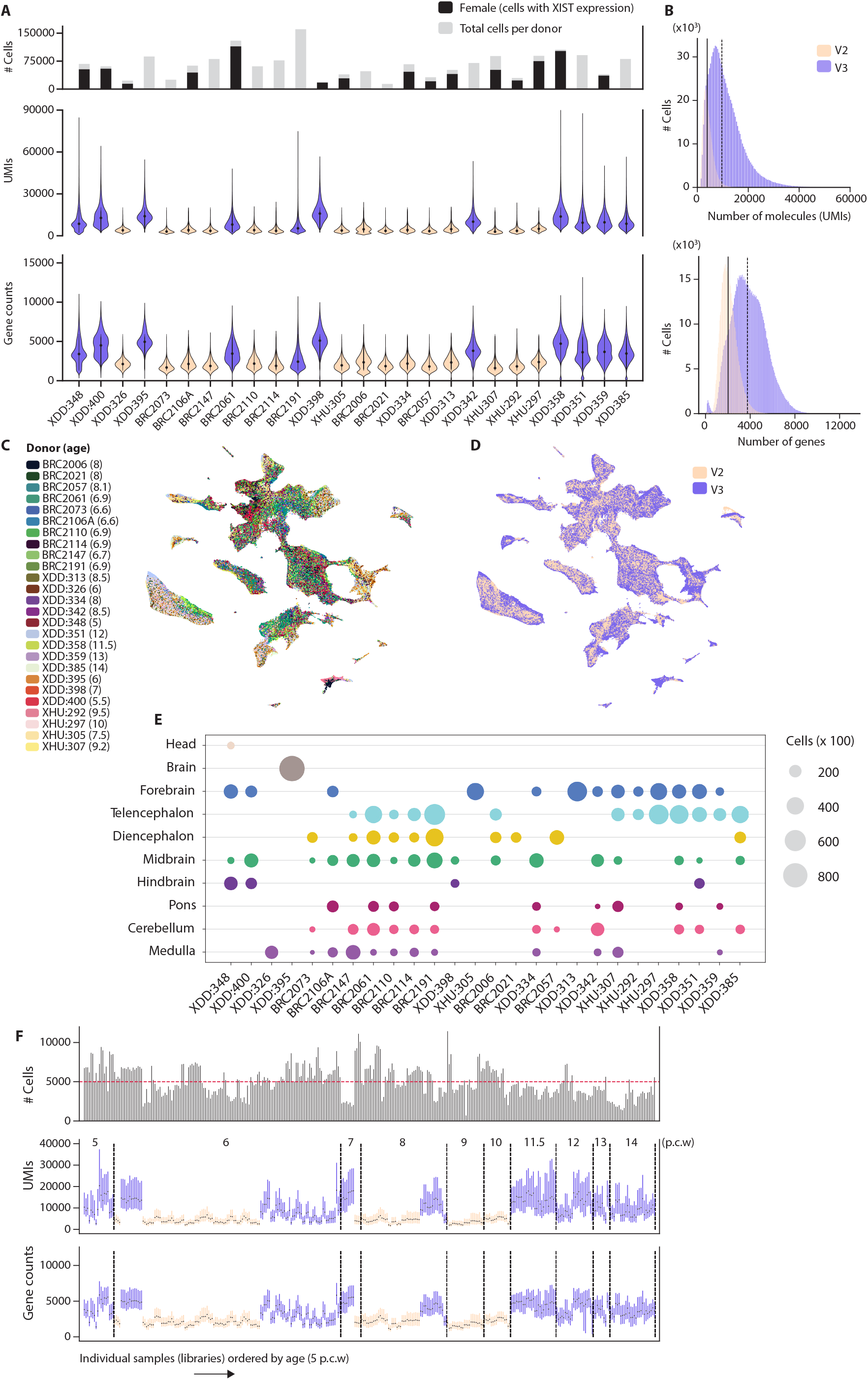
Donor metadata and sampling overview. **(A)** Violin plots show distribution of UMI (molecule) and gene counts per cell and donor, colors indicate chemistry version: V2 (peach) and V3 (violet). Bar chart (top) showing female donors (black) defined by the fraction of cells expressing the XIST gene. **(B)** Distribution of molecule counts (V2; median: 3 865, V3; median: 9 551) and gene counts (V2; median: 2 018, V3; median: 3 737) of all cells in the whole dataset. **(C)** tSNE embedding of whole dataset colored by donor, legend displaying all donors ang age per donor. **(D)** Whole dataset colored by chemistry. **(E)** Sampling overview per donor: dot size indicating number of cells sampled per region for each donor. **(F)** Vertical lines (10 to 90 percentile; dots: median) of UMI and gene counts distribution of individual sequencing libraries, sorted in ascending age order, dashed lines dividing beginning and end of whole timepoints. Bar chart (top) showing number of cells per library, mean shown in dashed line (red).

**Supplementary Figure 2.**
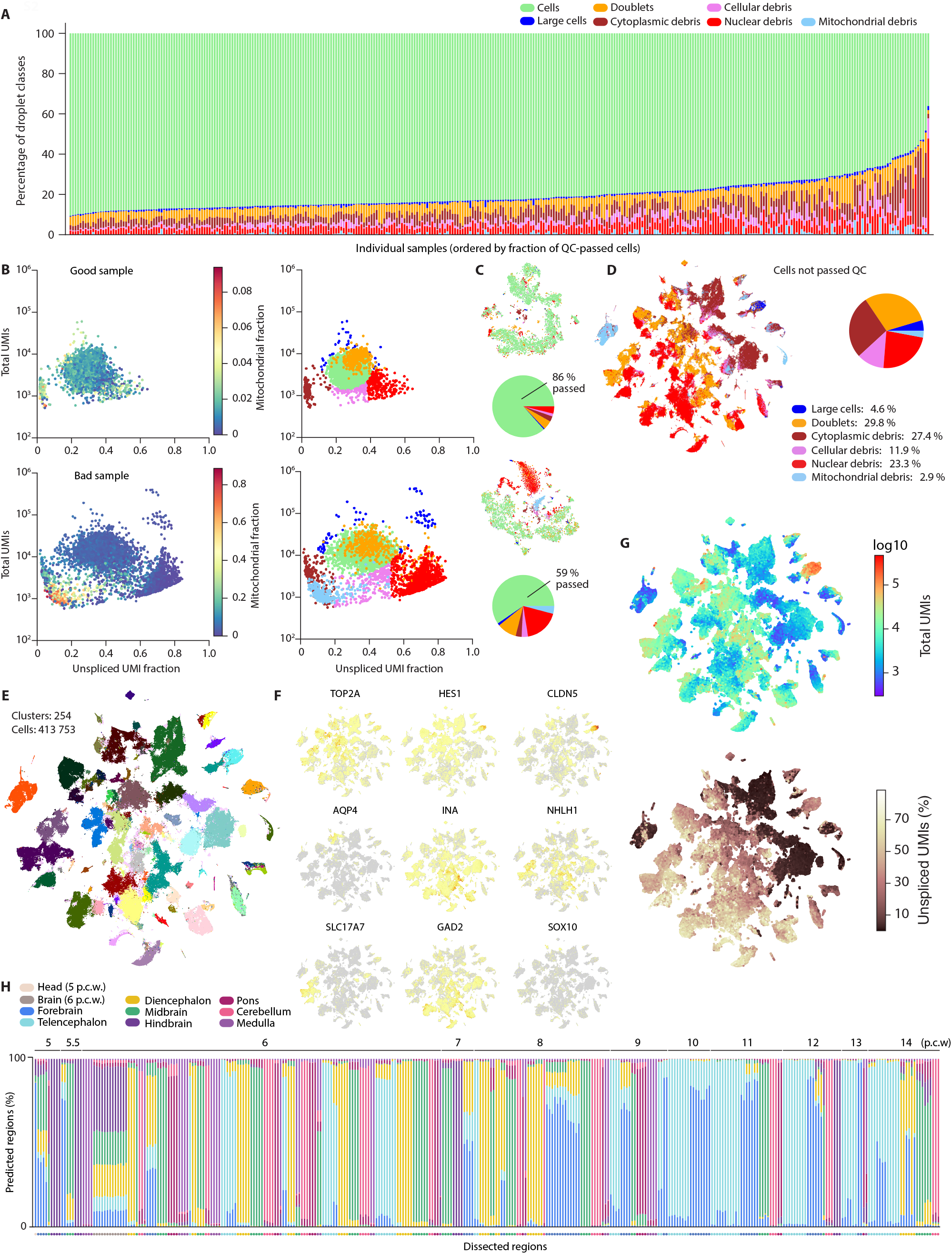
Quality control metadata and discarded cells. **(A)** Fraction of droplet class per individual sample/library, sorted in descending order of fraction of good cells. **(B)** Example of quality metrics in a good (top) and bad sample (bottom). Scatter plots (each dot is a cell/classified droplet) show mitochondrial fraction (left) and droplet class (right) defined by their unspliced UMI ratio (x-axis) and total UMIs (y-axis). Good sample: female donor XHU:297, sample ID 10×101_5, forebrain, 10 p.c.w., v2 chemistry. Bad sample: male donor XDD:351, sample ID 10×187_5, 12 p.c.w., v3 chemistry. **(C)** tSNE embeddings of the same samples (good: top; bad: bottom) after preliminary clustering to detect abundance of certain droplet groups. Pie charts showing the fractions of droplet classes of the cells in each sample. **(D)** Embedding of clusters of all cells that did not pass QC (discarded droplets) colored by droplet class. Pie chart showing percentages of droplet classes/cells that got discarded. **(E)** Clusters of discarded cells, colored by individual cluster (cell type). **(F)** Gene expression of typical markers: *TOP2A* (cycling), *HES1* (radial glia), *CLDN5* (endothelial), *AQP4* (pre-astrocytic), *INA* (neuronal, *NHLH1* (neuroblast), *SLC17A7* (excitatory neurons), *GAD2* (inhibitory neurons), *SOX10* (oligodendrocyte precursors). **(G)** Total molecules of bad droplets (top), fraction of unspliced molecules (bottom). **(H)** Predicted regions of each sequenced library. Each bar show fraction of cells predicted as a certain region per sample. Dots on x-axis indicate dissected region labels. Samples ordered in ascending age order.

**Supplementary Figure 3.**
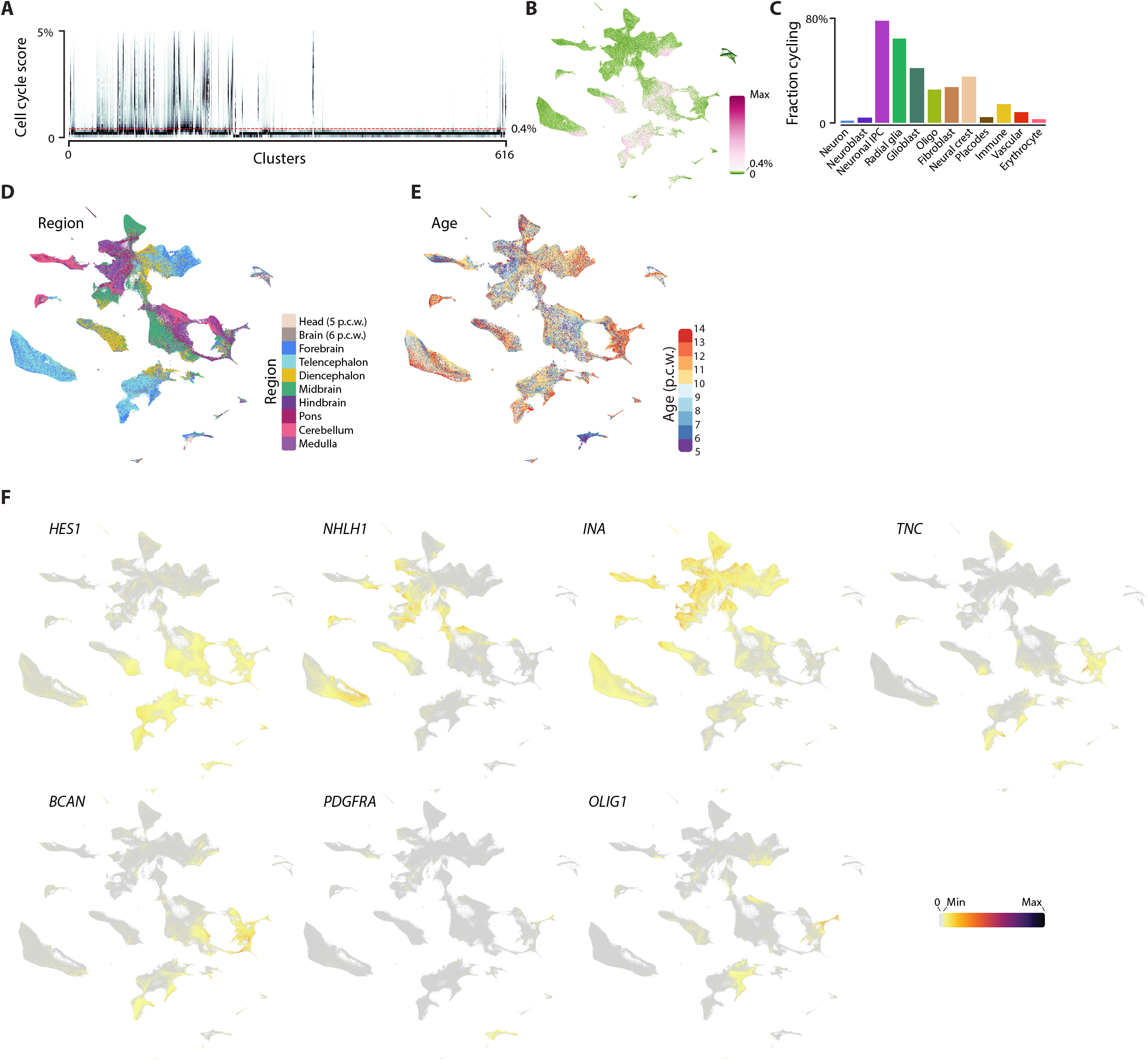
Metadata corresponding to Figure 1. **(A)** Cell cycle score shown as vertical histograms, one per cluster. **(B)** Cell cycle score shown on the tSNE. **(C)** Fraction of cells with cell cycle score > 0.4%, per cell class. **(D)** Region annotation shown on tSNE embedding. **(E)** Age shown on tSNE embedding. **(F)** Key markers used to identify major classes of cells.

**Supplementary Figure 4.**
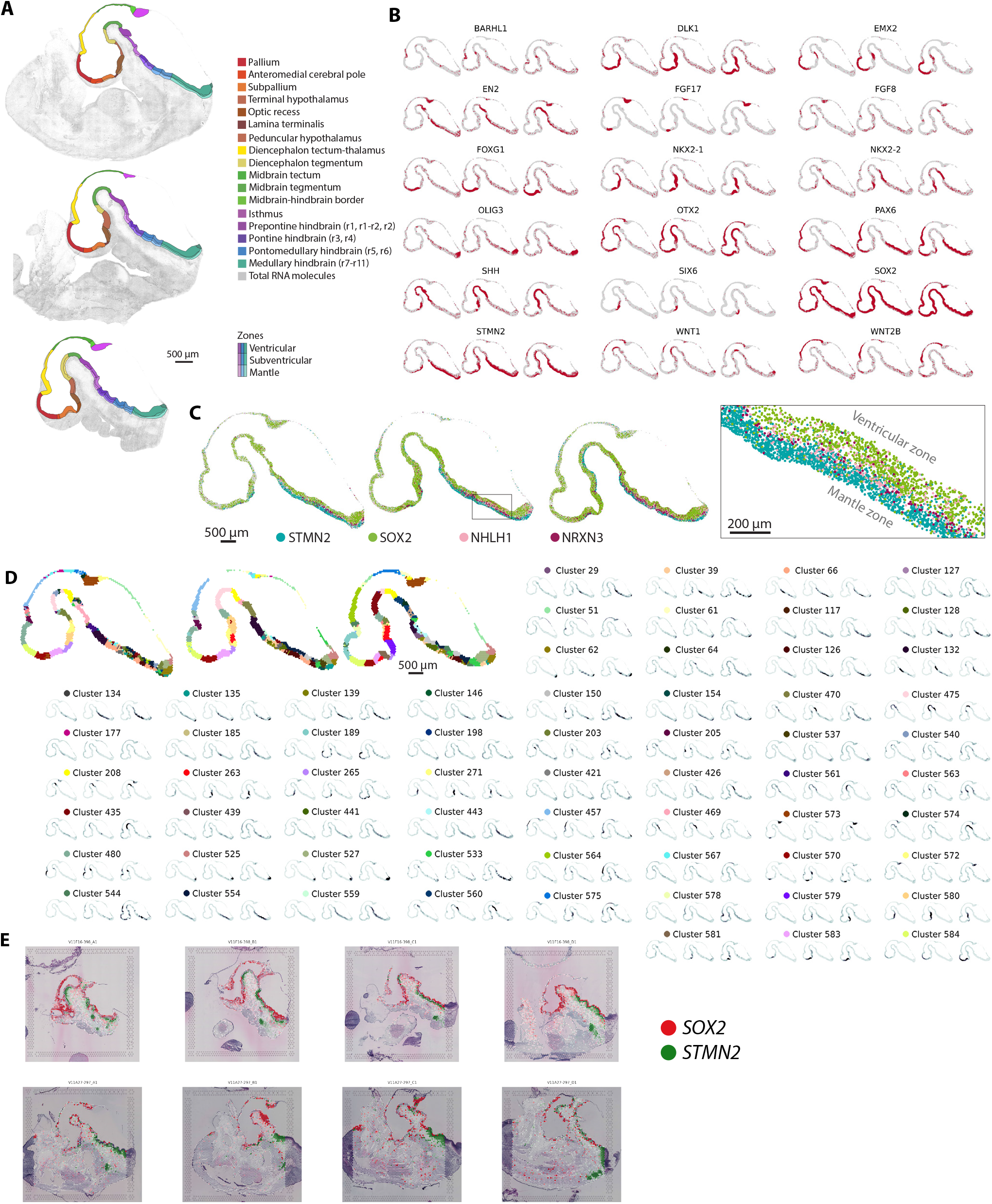
Spatial transcriptomics on 5 p.c.w. human brain. **(A)** Manually curated anatomical regions according to prosomeric model. **(B)** Expression of selected marker genes measured by multiplex RNA FISH. **(C)** Expression of genes marking radial layers: ventricular, subventricular and mantle zone. **(D)** Maximum posterior probability cluster assignments (top left) and individual cluster probability distributions indicating spatial locations of single-cell clusters. **(E)** Spatial transcriptomics (Visium) showing expression of STMN2 (neuorns) and SOX2 (radial glia) on four sagittal sections of the same embryo as in (A).

**Supplementary Figure 5.**
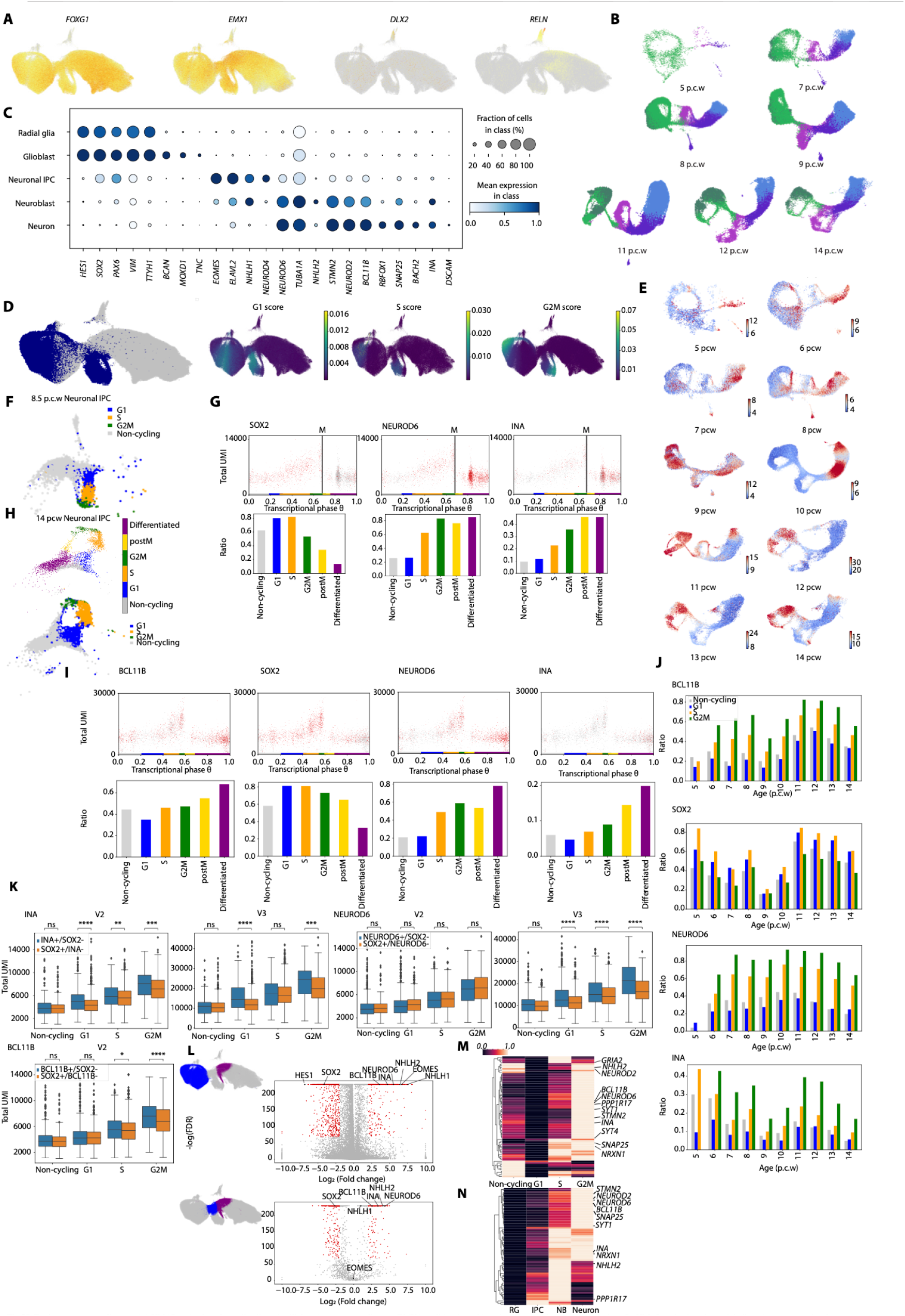
Excitatory neuronal lineage. **(A)** UMAP projections from all collected pallial excitatory neuron lineage (*EMX1*+ cells) coloured by selected patterning genes expression. **(B)** UMAP projections for each indicated post-conceptional ages pallium cells (*EMX1*+ cells) coloured by major cell classes. **(C)** Marker genes expression for major cell classes. Dotplot showing mean expression (color) and ratio of cells expressing the gene in each major cell class (circle size). **(D)** UMAP projections from all collected pallial excitatory neuron lineage (*EMX1*+ cells) colored by cell cycle gene expression (left) and colored by cell cycle phase score (G1, S, G2M) (right). RNA-velocity length predicted by scvelo for 5-14 p.c.w. One donor was selected for each p.c.w. UMAP projection of cortical *EOMES*+ cells in 8.5 p.c.w colored by progenitor states inferred by cell cycle phase score. Cortical *EOMES*+ cells from 8.5 p.c.w UMI counts per cell as a function of the transcriptional phase show growth trend followed by a drop in RNA counts (UMI) that identifies mitosis (marked by M). Cells expressing *SOX2, NEUROD6*, and *INA* are marked in red (top). Bar plots show the ratio of cells in each progenitor state expressing *SOX2, NEUROD6*, and *INA* (bottom). **(H)** UMAP projection of cortical *EOMES*+ cells from 14 p.c.w colored by progenitor states inferred by cell cycle phase score. **(I)** *EOMES*+ cells from 14 p.c.w UMI counts per cell as a function of the transcriptional phase show growth trend followed by a drop in RNA counts (UMI) that identifies mitosis (marked by M). Cells expressing *BCL11B, SOX2, NEUROD6*, and *INA* are marked in red (top). Bar plots show the ratio of cells in each progenitor state expressing *BCL11B, SOX2, NEUROD6*, and *INA* (bottom). **(J)** Bar plots show the ratio of IPCs expressing *BCL11B, SOX2, NEUROD6*, and *INA* in each progenitor state for 5-14 p.c.w samples. **(K)** Box plots showing RNA counts in *INA+/SOX2-*vs. *SOX2+/INA-*IPCs by progenitor states in Chromium version 2 and 3 samples as indicated (top left). Box plots showing UMI counts in *NEUROD6+/SOX2-*vs. *SOX2+/NEUROD6-*IPCs by progenitor states in Chromium version 2 and 3 samples (top right). Box plot showing UMI counts in *BCL11B+/SOX2-*vs. *SOX2+/BCL11B-*IPCs by progenitor states in Chromium version 2 samples (bottom left). **(L)** Select genes are annotated on a volcano plot illustrating differential expression between radial glia and differentiating neuronal IPCs (top). Significant differential expressed genes are coloured red. Miniature UMAP marks the group of cells that were compared (blue – radial glia cells, purple – differnitating IPCs). Select genes are annotated on a volcano plot illustrating differential expression between IPCs entering the cell cycle and differentiating IPCs using the genes selected in the RG vs. differentiating IPCs test (bottom). Significant differential expressed genes are coloured red. Miniature UMAP marks the group of cells that were compared (blue – IPCs entering the cell cycle, purple – differnitating IPCs). **(M)** Expression heatmaps of genes associated with neuronal differentiation in different progenitors state. **(N)** Expression heatmaps of genes associated with neuronal differentiation in different classes (RG - radial glia, IPC-intermediate progenitor cell, NB-neuroblast).

**Supplementary Figure 6.**
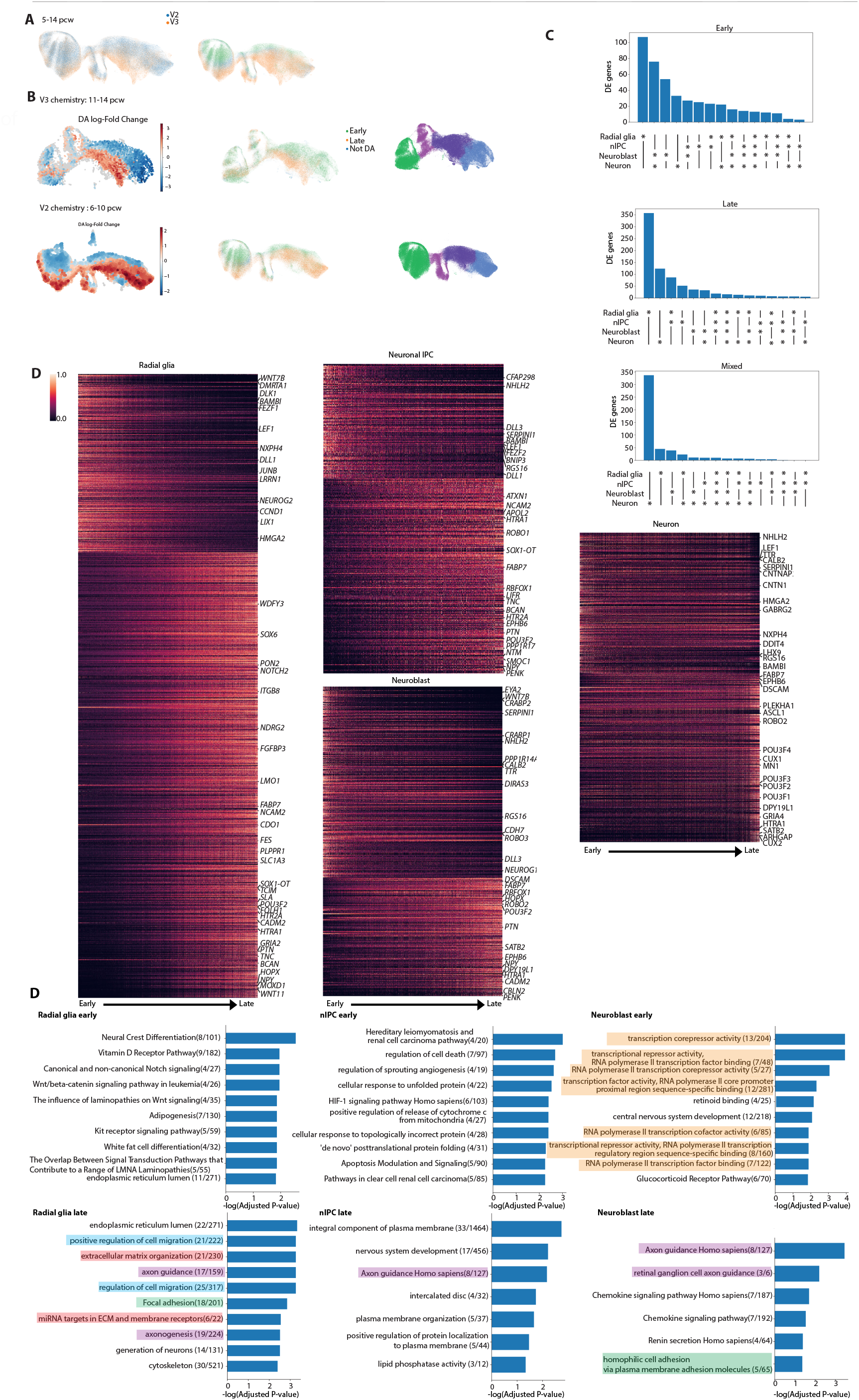
Excitatory neuronal lineage – early vs. late states. **(A)** UMAP projection all collected cells of the pallial excitatory neuronal lineage (*EMX1*+ cells) colored by Chromium versions (Left) and by the subsets used for differential expression test (Right). **(B)** Differential abundance plot of Chromium version 3 samples from 11-14 p.c.w of the pallial excitatory neuron lineage (top left) and Chromium version 2 samples from 6-10 p.c.w in pallial excitatory neuron lineage (bottom left). Each point represents a neighborhood, the size of points is proportional to the number of cells in the neighborhood. Neighborhoods are colored by their log-fold change in abundance between early and late post-conceptional age. Neighborhoods showing significant enrichment (SpatialFDR<10%) are colored. Neighborhoods with higher abundance of cells from early ages are colored blue and neighborhoods with higher abundance of cells from later ages are colored red. UMAP projections colored by subset used for differential expression test and major cell classes for Chromium version 3 samples from 11-14 p.c.w (top panel middle and right, respectively), Chromium version 2 samples from 6-10 p.c.w (bottom panel middle and right, respectively). **(C)** Bar plot showing the number of significant genes associated with early (top), late (middle) and more than one type of change (“mixed”, bottom) in each major cell class. **(D)** Expression heatmaps of differential expressed genes in radial glia cells, IPCs, neurobalsts and neurons. The mean expression for each neighborhood defined by milo is shown. The neighborhoods are ordered according to Log(fold change) of the differential abundance between early and late states. Selected gene names are shown. **(E)** Gene ontology and pathway enrichment analysis results. Showing the lowest adjusted p-value results. In case of duplicated terms only the one with the lower FDR is shown. Axon related gene ontologies are colored purple, cell adhesion related GO are colored green, transcription related terms are colored orange, extra cellular matrix related GO are colored red, and cell migration related GO are colored blue.

**Supplementary Figure 7.**
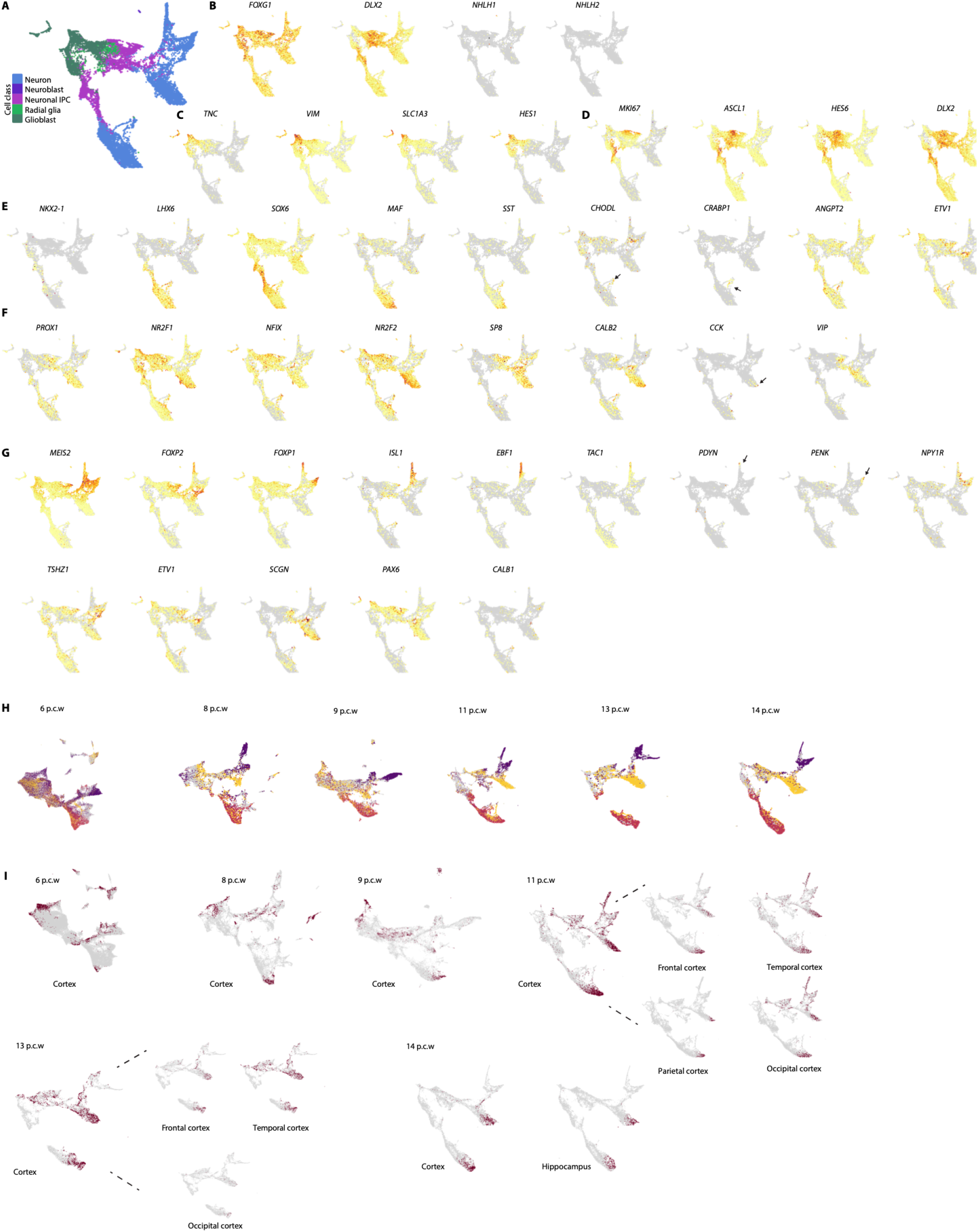
Forebrain GABAergic neurons – cortical migration. **(A-G)** UMAP projections of cells from 12 p.c.w forebrain GABAergic lineage colored by **(A)** major cell classes **(B)** expression of selected patterning genes **(C)** expression of radial glia marker genes **(b)** expression of IPCs marker genes **(E)** expression of MGE and derived neuronal types marker genes **(F)** expression of CGE and derived neuronal types marker genes **(G)** expression of LGE and derived neuronal types marker genes. Only *EMX1*-cells are shown **(H)** UMAP projections of cells from different p.c.w colored by GE marker genes score. **(I)** UMAP projection of cells from different p.c.w colored by the indicated Subregion (brown). In case of several cortical dissections for a time point, showing all the samples together and zoom in figures for each dissection.

**Supplementary Figure 8.**
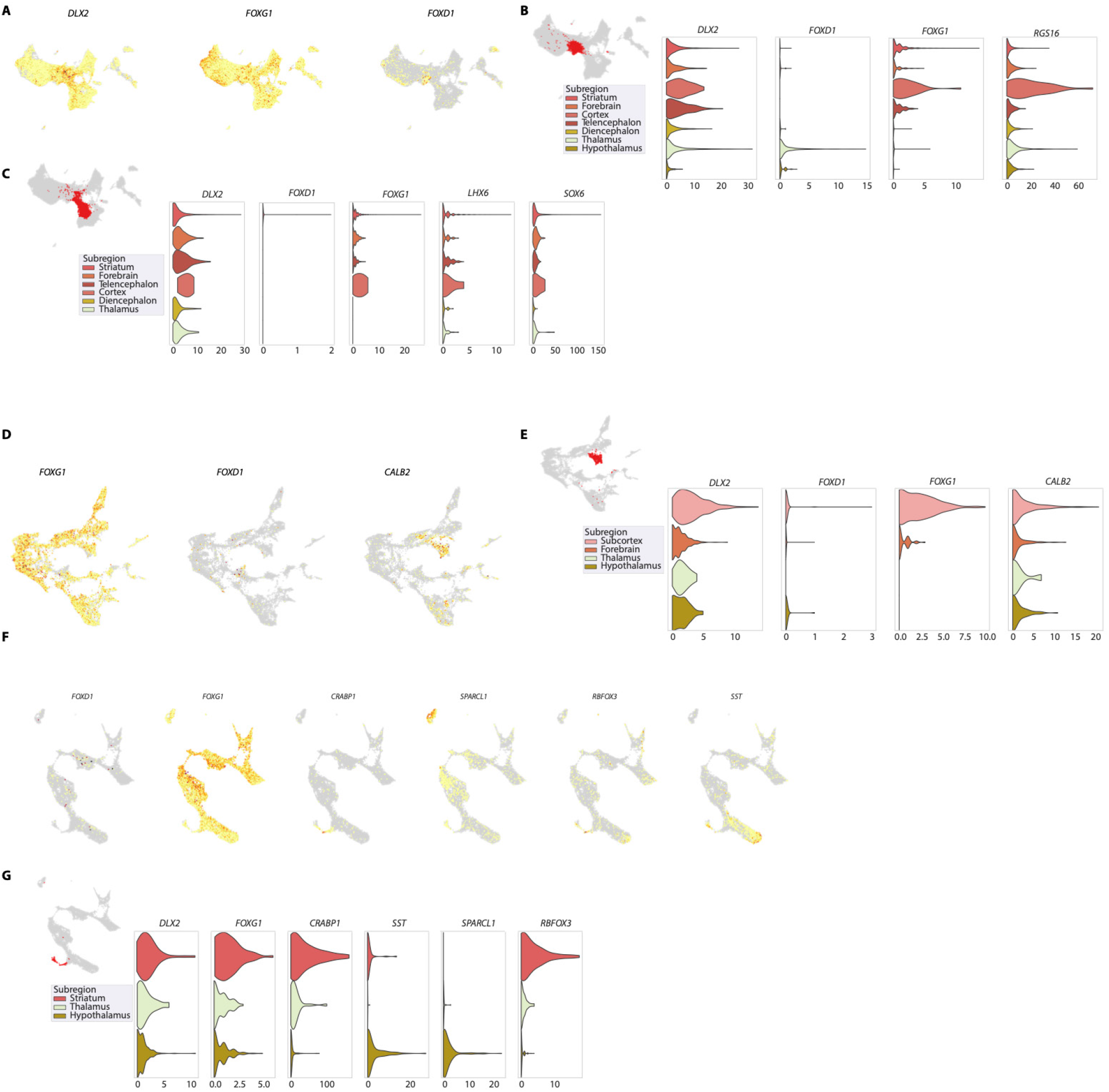
*DLX2*+ cells in the thalamus and hypothalamus. **(A)** UMAP projection of 6 p.c.w forbrain GABAergic lineage cells colored by expression of selected genes **(B)** Violine plots showing expression of *DLX2, FOXD1, FOXG1* and *RGS16* in cluster 9 of 6 p.c.w lineage in different subregions. Miniature showing cluster 9 cells (red). **(C)** Violine plots showing expression of *DLX2, FOXD1, FOXG1, LHX6*, and *SOX6* in cluster 14 of 6 p.c.w lineage in different subregions. Miniature showing cluster 14 cells (red). **(D)** UMAP projection of cells from 8 p.c.w forbrain GABAergic lineage colored by expression of selected genes. **(E)** Violine plots showing expression of *DLX2, FOXD1, FOXG1* and *CALB2* in cluster 15 of 8 p.c.w lineage in different subregions. Miniature showing cluster 15 cells (red) **(F)** UMAP projection of cells from 14 p.c.w forbrain GABAergic lineage colored by expression of selected genes. **(G)** Violine plots showing expression of *DLX2, FOXG1, CRABP1, SST, SPARCL1*, and *RBFOX3* in cluster 0 of 14 p.c.w lineage in different subregions. Miniature showing cluster 0 cells (red).

**Supplementary Figure 9.**
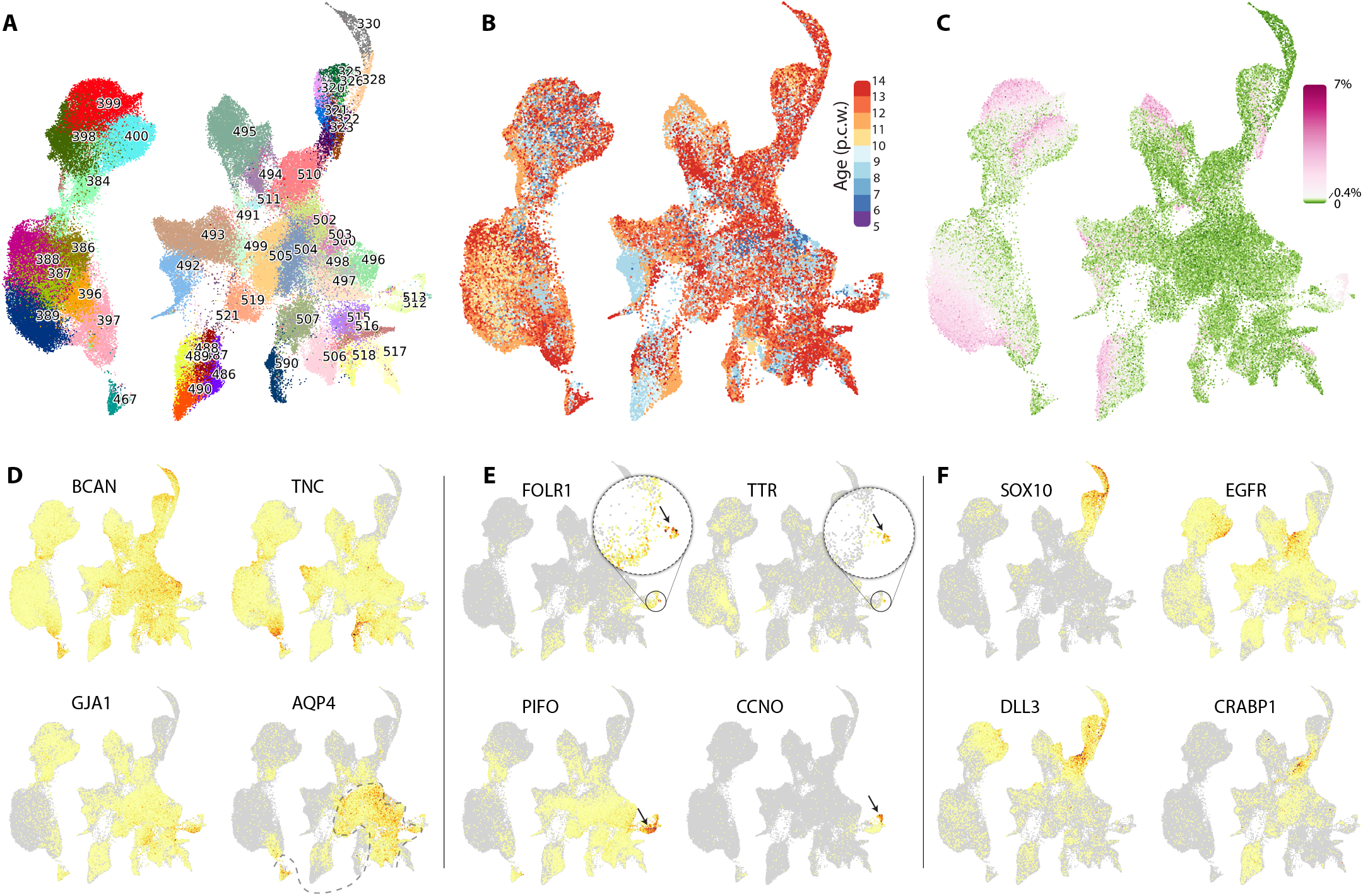
Glioblasts. **(A)** Clusters shown on tSNE embedding. **(B)** tSNE colored by age. **(C)** tSNE colored by cell cycle gene expression. **(D)** Expression of the indicated genes shown on the tSNE embedding.

**Supplementary Figure 10.**
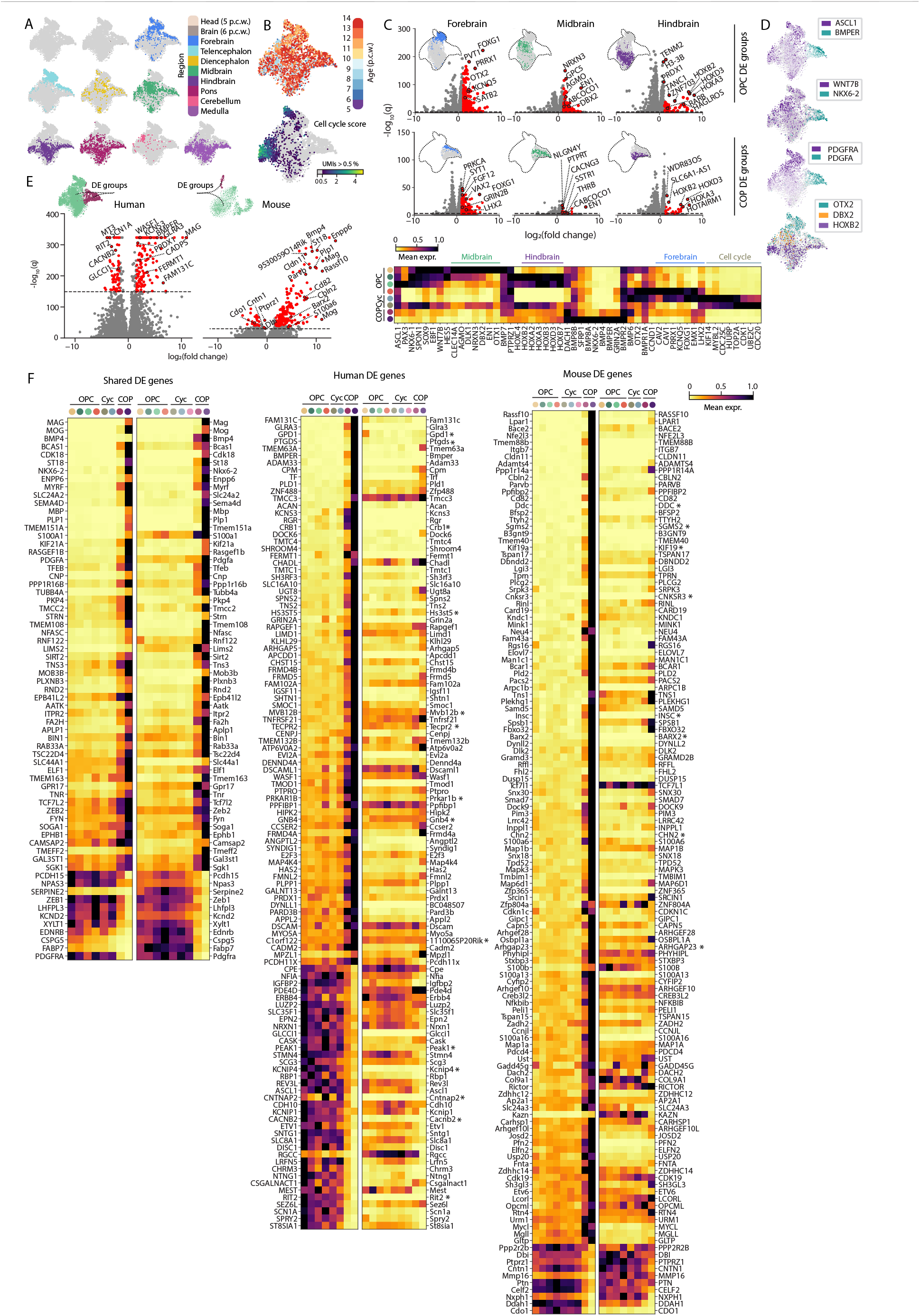
Regionalization in OPCs and species comparison. **(A)** tSNE embeddings labeled by individual dissected regions. **(B)** Embeddings of displaying age metadata (top) and cell that are cycling (defined by cells that have > 0.5 % molecules from cell cycle genes; bottom). **(C)** Volcano plots of differentially expressed genes in fore-, mid- and hindbrain of OPCs (top) and COPs (bottom). Miniature tSNEs showing test groups within each region for the differential gene expression analysis. Heatmap showing region-specific genes of human OPCs, cycling OPCs and COPs after hierarchical clustering. **(D)** Embeddings displaying expression OPC- and COP-specific genes (top three) and patterning genes: *OTX2* (forebrain), *DBX2* (midbrain) and *HOXB2* (hindbrain) at bottom. **(E)** Volcano plots of differentially expressed genes between OPCs and COPs in human and mouse respectively. Miniature tSNEs show test groups (green: OPCs, plum: COPs). **(F)** Heatmaps showing gene expression of shared genes in mouse and human (right), human- (middle) and mouse-specific genes (right).

